# Histone H3 cysteines as redox switches in the nucleosome: molecular modeling of S-sulfenylation and its structural consequences

**DOI:** 10.1101/2024.10.07.616940

**Authors:** Samuel D. Mador, Roy González-Alemán, Yuya Qiu, Francisco Javier Patiño-Lopéz, Yani Kedjar, Mailys Duch, Yasaman Karami, Emmanuelle Bignon

**Affiliations:** Université de Lorraine, CNRS, UMR 7019 LPCT, F-54000 Nancy, France; Université de Lorraine, Inria, F-54000 Nancy, France

**Keywords:** Nucleosome, Histone oxidation, *S*-sulfenylation, Hydrogen peroxide, Redox signaling, Molecular dynamics

## Abstract

Oxidation of buried histone cysteines links cellular redox signaling to chromatin regulation. Histone H3 carries two such residues: C96, unique to the H3.1 variant, and C110, shared by most H3 variants. Oxidation of H3.1 C96 by H_2_O_2_ drives histone variant exchange and chromatin opening in cancer cells, and C110 undergoes *S*-glutathionylation, yet both thiols are shielded from the solvent. How they are oxidized and what the modification does to the nucleosome structure remain unknown. Combining all-atom molecular dynamics, QM/MM and free energy calculations, we reveal *S*-sulfenylation mechanisms at both cysteines within the nucleosome core particle, from oxidant access to its structural consequences. Reaching either cysteine costs ~12 kcal/mol, paid mostly for the desolvation of H_2_O_2_ independently of the entry route. The reaction is then enabled in the hydrophobic environment by the presence of a single neighboring base. Such a residue is already in place at C110 (H3D123), but must first reorganize at C96 (H3E97), with barriers of ~18 kcal/mol at both sites. Once formed, the modification acts well beyond its site. C96 *S*-sulfenylation acts on DNA-histone interfaces, on elements recognized by chromatin remodelers and chaperones (H2AE91 at the acidic patch, H2A C-term release), and on the H4K91-H2B′D68 salt bridge bridging the two halves of the nucleosome. C110 oxidation mainly favors unwrapping and DNA destabilization at the dyad. These results provide a structural framework for understanding histone-mediated redox signaling and its implication in chromatin remodeling.

**Highlights:**

- The histone core shields its cysteines from H_2_O_2_ behind a desolvation barrier
- A single neighboring base enables cysteine oxidation in a hydrophobic environment
- H3.1 oxidation modulates remodeler-binding sites to prime histone variant exchange
- C96 and C110 oxidation leave distinct dynamic signatures in the nucleosome

## 1. Introduction

DNA compaction is a dynamic phenomenon regulated by different factors. Among them, post-translational modifications (PTMs) of histone proteins are crucial, and hundreds of them have been characterized to date [1]. Histone PTMs located on the histone tails can directly affect tail-DNA electrostatic interactions and can trigger the binding of partner proteins [2]. Besides, PTMs located on the histone core can promote either nucleosome disassembly when located near the dyad, or DNA unwrapping when positioned on the lateral surface of the histone core [3]. Nevertheless, structural data remain scarce, limiting our understanding of these regulatory mechanisms at the atomic scale.

This lack of data is especially marked for oxidative PTMs. In the nucleosome context, histone H3 carries two cysteine residues that are targets of redox chemistry: C96, found exclusively on the H3.1 variant, and C110, conserved across most H3 variants [4] - Figure 1. H3C96 is buried at the H3/H4 interface and sits in a much more hydrophobic environment than C110, yet it has emerged as a central redox switch [5]. Indeed, Bonini and coworkers have shown that H3.1 behaves as a chromatin-embedded redox sensor, whose oxidation is directly modulated by nuclear and mitochondrial reactive oxygen species [6]. Specifically, oxidation of H3.1 Cys96 by H_2_O_2_ might destabilize the nucleosome, promoting H3.1 eviction and its replacement by the H3.3 variant. In breast cancer cells, this mechanism was shown to favor cell plasticity, metastasis, and multidrug resistance, while scavenging nuclear H_2_O_2_ or mutating the cysteine back to a serine (H3.1 C96S, as found in H3.2) suppressed this plasticity and restored chemosensitivity [6]. On the other hand, the importance of H3C110 in redox-related events was strikingly illustrated by Kurdistani and coworkers, who showed that the (H3-H4)_2_ tetramer itself behaves as a copper reductase enzyme, catalyzing the reduction of cupric (Cu^2+^) to cuprous (Cu^1+^) ions at the H3-H3′ interface bearing this cysteine [7]. This activity has been linked to cellular copper homeostasis, mitochondrial Fe-S cluster metabolism, and disease [8, 9], positioning C110 as a catalytic redox center within the nucleosome in addition to its known role as a site of regulatory oxidation. Indeed, H3C110 is also known to undergo different types of oxidative modifications, including *S*-sulfenylation, *S*-sulfonylation and *S*-glutathionylation [10, 11]. Experimental studies have shown its *S*-glutathionylation promotes chromatin opening [12] and is linked to cell proliferation [13], while some of our recent computational work has underlined the destabilization of nucleosome structure by its hyperoxidation [14].

**Figure 1:**
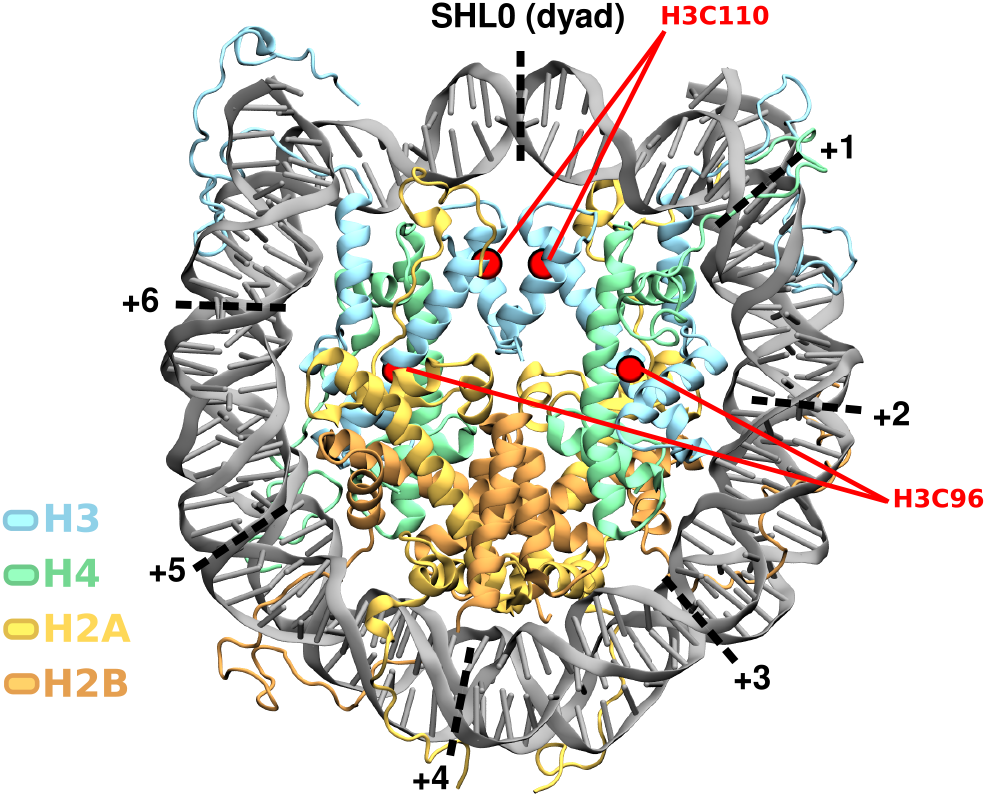
Structure of the nucleosome core particle containing the H3.1 variant. Histones H3, H4, H2A, and H2B are colored blue, green, yellow, and orange, respectively. The redox-active cysteines H3C110 and H3C96 are highlighted in red, and superhelical locations (SHLs) are indicated along the DNA.

Taken together, these findings show that both nucleosomal cysteines, H3C96 and H3C110, sit at the crossroads of redox biology and chromatin regulation, yet no structural or atomistic picture currently exists of how oxidation of either residue can occur and perturb the nucleosome architecture. While computational studies of nucleosome dynamics and the effects of some mainstream PTMs have been reported (reviewed in [15]), oxidative histone modifications remain mostly unexplored from a theoretical point of view.

In this work, our aim is to address this lack of data by investigating the oxidation of both cysteines of histone H3 within the nucleosome core particle (NCP), focusing primarily on *S*-sulfenylation. Because both residues are buried within the histone core and largely shielded from the solvent, we specifically investigate: *i)* how H_2_O_2_ can access these cysteines, *ii)* how the *S*-sulfenylation reaction can take place in such a hydrophobic environment, and *iii)* how the resulting modification subsequently impacts the nucleosome structure and histone-DNA interactions. To address these questions, we employed a multiscale approach combining unbiased classical and QM/MM-MD simulations, enhanced sampling methods, and free energy calculations. This combination allowed us to capture both the reactive step (i.e., the access of H_2_O_2_ to the buried cysteine and the ensuing chemistry) and the structural consequences at the nucleosome scale. We additionally performed simulations on *S*-nitrosylated H3C110 for comparison, given its role as reaction intermediate toward *S*-glutathionylation – a modification known to occur on this residue.

We reveal that: *i)* H_2_O_2_ accesses C96/C110 via specific gated pathways, *ii)* the *S*-sulfenylation reaction can proceed in the hydrophobic environment with the assistance of a nearby base, and *iii)* oxidative modifications perturb the nucleosome dynamics by local and allosteric effects. These results provide an atomistic picture of how oxidation of these two cancer- and redox-relevant cysteines perturbs the NCP dynamics, rationalizing the cellular and biochemical findings reported by Bonini and coworkers for H3.1 C96, while also shedding light on the so far unexplored structural consequences of H3.2 C110 oxidation.

## 2. Results

### 2.1. H_2_O_2_ access pathways to C96 and C110

Cysteines C96 and C110 are deeply buried at the H3–H4 and H3–H3′ interfaces, respectively. While it is known that these residues act as redox centers in the nucleosome, how oxidative species access these sites for subsequent modification remains unclear. Therefore, we started by probing the mechanisms driving H_2_O_2_ binding to both sites in the NCP with the H3.1 variant bearing both cysteines.

#### H_2_O_2_ approaches both cysteines but never reaches them through simple diffusion

To assess the possibility of spontaneous binding of H_2_O_2_ to the cysteine sites, *i*.*e*., through a simple diffusion process, we conducted unbiased MD simulations of the nucleosome in the presence of H_2_O_2_ molecules in solvent. Along the simulations, H_2_O_2_ is found within 10 Å of C96 and C110 for about 29.8% and 28.1% of the simulation time, respectively, and it can get as close as 5.75 Å of C96 and 5.34 Å of C110 (Figure 2A). Near H3C96, H_2_O_2_ spontaneously binds to a pocket formed by H4L90, H4R95, H4F61, H3A95, and H3Y99, with a maximum persistency of ~38 ns (Figure 2B and S1). In the vicinity of H3C110/H3′C110 on the other hand, H_2_O_2_ binds to a pocket formed by H3D106, H3L109, H3R129, H3I130, and H3R131, with a maximum residency time of ~43 ns (Figures 2C and S1). MDpocket analysis of persistent cavities along MD simulations of the NCP with both variants identifies the binding site near H3.1C96 and H3.2S96 (Figure S2A,C). Around C110, the closest persistent pocket is found at the adjacent position (close to H3L109 and H3’L126), while two other pockets are found above and below the C110 copies (Figure S2B,D), the latter corresponding to the binding site described above.

**Figure 2:**
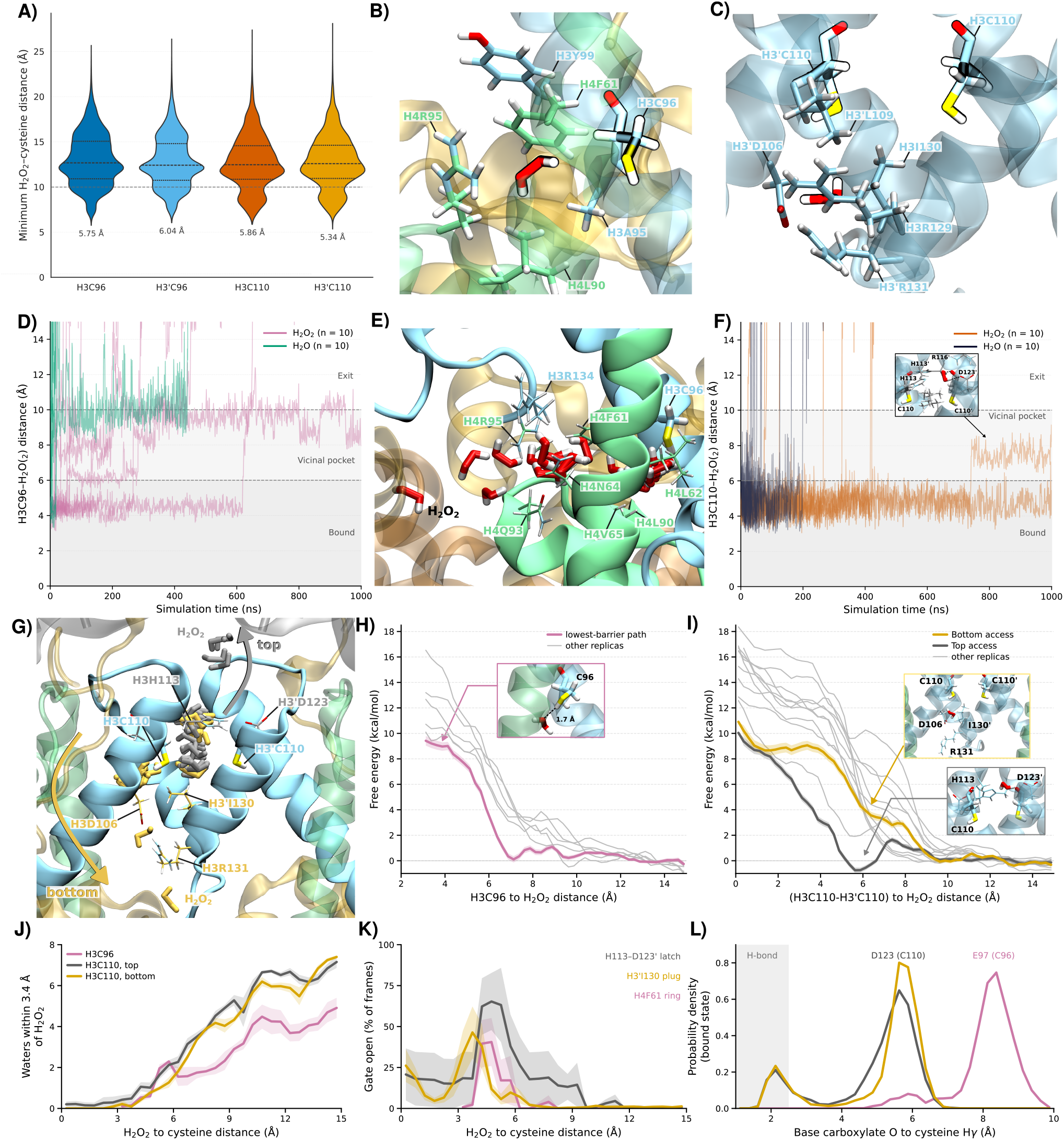
Molecular basis of H_2_O_2_ access to the H3.1 C96 and C110 cysteine sites within the nucleosome core particle. (A) Distribution of the H_2_O_2_–cysteine minimum distance in unbiased MD simulations of the nucleosome core particle with 0.15 M H_2_O_2_ in the bulk. (B,C) Conformations at the minimum distance observed in (A) for H3C96 (B) and H3C110 (C). (D,F) Residence of H_2_O_2_ and H_2_O near H3C96 (D) and H3C110 (F) along the post-steered MD relaxation replicas; the inset in (F) shows H_2_O_2_ in the top vicinal pocket. (E,G) H_2_O_2_ exit pathway from the H3C96-bound (E) and H3C110-bound (G) states. For H3C110, H_2_O_2_ leaves either via the top (grey; gate H3H113–H3′D123) or the bottom (yellow; gate H3D106, H3′I130, H3R131) of the site. (H,I) Umbrella sampling free energy profiles for H_2_O_2_ access to H3C96 (H) and H3C110 (I; top grey, bottom yellow). Six replicas per pathway; the lowest-barrier one is in color (shade: WHAM bootstrap error), the others in grey. Each profile is zeroed on its own bulk plateau. Insets show conformations along the paths. (J) Waters within 3.4 Å of H_2_O_2_ along the three routes. (K) Fraction of frames in which the gate of each route is open, *i*.*e*. outside the range it samples in bulk: H4F61 ring flipped (C96), H3H113–H3′D123 latch (C110, top) and H3′I130 plug (C110, bottom). (L) Distance between the carboxylate oxygens of the candidate base (H3D123 for C110, H3E97 for C96) and the cysteine Hγ in the bound state; the shaded band marks H-bonding distance.

More globally, H_2_O_2_ binds DNA and histone residues in similar proportions (Figure S3). On the histone core, the main persistent interactions are observed at the histone–DNA interface at SHL±0.5 and SHL±1.5 (Figure S4). There, the H_2_O_2_ molecules can get trapped between H4 R39/K44/R45, H3 P43/G44 and the DNA minor groove at SHL±0.5, for up to 605 consecutive ns. The second main site is H3 K64/L65 at the N-term end of helix α1, facing DNA at SHL±1.5, with stays of up to 291 ns. Both sites are symmetric and found in all replicas. Shorter-lived binding sites (tens of ns) are found at buried protein-protein interfaces: the H4 α2 / H2B α3 bundle (H4 R67/D68 with H2B R99/L100, near the H3C96 binding site described above), and H3 E94 against the H2A′ docking domain (Q104–G106). Of note, several of these sites are located next to redox-active tyrosine residues reported to be nitrated in cells, namely H3 Y41 at the SHL±0.5 site and H4 Y72 in the H4 α2 / H2B α3 bundle [10, 11].

Although H_2_O_2_ spontaneously approaches both H3C96 and H3C110 sites by binding at specific nearby pockets, the complete binding to either residue was never observed, suggesting that an energy barrier prevents the oxidant from reaching these residues by simple diffusion.

#### H_2_O_2_ shape enables stabilizing interactions at both C96 and C110

To investigate the access and stability of H_2_O_2_ once bound to the buried cysteines, we performed steered MD simulations to force H_2_O_2_ into the vicinity of both C96 and C110. We then relaxed the resulting bound state through unbiased MD simulations to probe the residence time at each site and the possible exit pathways. Similar simulations were performed with H_2_O to assess the specificity of H_2_O_2_ binding in these hydrophobic environments.

At C96, H_2_O_2_ can remain bound for up to 600 ns (7/10 replicas > 50 ns), yet systematically drifts away from the cysteine (Figure 2D). In its bound state, it interacts mainly with backbone atoms of the hydrophobic residues H4L62, H4V65, H4L90, and with the cysteine itself (Figure S2E). H_2_O_2_ systematically exits through the previously described vicinal binding pocket, that seems to act as an antechamber in which H_2_O_2_ can remain for several hundred nanoseconds. In addition to the residues identified above, H3R134, H4N64, and H4Q93 are also found to interact with H_2_O_2_ along its exit pathway (Figure 2E). Interestingly, H_2_O does not populate this antechamber, and consistently exits the C96 vicinity within 10–20 ns of simulation time (Figure 2D). The higher persistence of H_2_O_2_ in the cavity appears to be linked to its more elongated shape, which allows for simultaneous interactions with nearby polar backbone atoms and with C96 within the otherwise hydrophobic pocket.

At C110, H_2_O_2_ can remain bound for as long as 1 µs (*i*.*e*. the maximum simulation time; 9/10 replicas > 50 ns), while H_2_O exits within 10–200 ns (5/10 replicas > 50 ns; Figure 2F). While its position in the cavity fluctuates, H_2_O_2_ is mostly stabilized by interacting with the backbone carbonyl oxygens of the two C110 copies, just below H3H113 and its symmetric counterpart (Figure S2F). Analysis of its complete spontaneous exit from the cavity points to two distinct exit routes: either through the top of the cavity, via a cryptic pocket formed between H3H113 and H3′D123 where it can get stabilized for hundreds of nanoseconds, or through the bottom of the cavity via the previously described vicinal pocket flanked by H3D106, H3′I130, and H3R131 (Figure 2G).

Together, H_2_O_2_ binding at both cysteines is reversible and specific to H_2_O_2_ over H_2_O, with a longer residence at C110 than at C96. H_2_O_2_ exit occurs through well-defined pathways, which raises the question of the energetic cost of the entry process.

#### The nucleosome core shields H3 cysteines from H_2_O_2_ through a desolvation barrier

While H_2_O_2_ can spontaneously exit the vicinity of the cysteines via specific pathways, its entry from the bulk solvent should be associated with a non-negligible energy barrier since no spontaneous complete binding could be observed. To quantify this barrier, we performed umbrella sampling along each of the three pathways described above, in six independent replicas per pathway (Figures 2H,I and S5). Along these paths, H_2_O_2_ makes reproducible contacts, observed in all six replicas, with H4R95, H4F61 and H3Y99 in the antechamber leading to C96, with H3′D106, H3′L109 and H3I130 in the bottom channel to C110, and with H3H113, H3′H113 and H3L126/H3′L126 around the C110 pocket on both C110 routes (Figure S6).

Access to C96 proceeds via a single pathway. In the lowestenergy replica, the bound state lies 9.2 kcal/mol above bulk and has no minimum, suggesting that it is metastable rather than thermodynamically favorable state, which is consistent with the spontaneous exit observed in the unbiased simulations described above (Figure 2H). Two shallow intermediates, 0.3–0.4 kcal/mol above bulk and at 9.9 and 7.4 Å from C96, correspond to H_2_O_2_ bound within the vicinal pocket described above (Figures 2B and S2), and the entry barrier itself is found between 7 and 4 Å. Importantly, pre-reactive structures are sampled in the bound state, with the distance between C96@HG and H2O2@O atoms as low as 1.69 Å.

The two pathways leading to C110 end at the same height, with 9.3 and 9.9 kcal/mol for the best top-entry and bottom-entry replicas, respectively, in both cases without a bound-state minimum (Figure 2I). Access via the top of the cavity proceeds through a stabilized intermediate, 0.7 kcal/mol below bulk at 5.9 Å and reached over a 1.5 kcal/mol activation barrier, with H_2_O_2_ just above the open H3H113–H3′D123 gate and the H3H113 side chain rotated outwards. Access via the bottom shows no such intermediate and climbs monotonically once H_2_O_2_ engages the H3D106/H3R131/H3′I130 gate. For both pathways the end states exhibit pre-reactive structures, with the H2O2@O to C110@SG distance reaching 2.78 Å (top pathway) and 2.73 Å (bottom pathway).

Averaged over the six replicas, the three routes are indistinguishable (11.7 ± 2.4, 14.0 ± 3.1 and 13.9 ± 2.4 kcal/mol free energy barrier for C96, top and bottom C110 entries respectively (Figure S5D)), which implies that, the cost of access is a property of the nucleosome core rather than of a particular route. Indeed, what is paid for is desolvation, as H_2_O_2_ loses its entire first hydration shell on the way in (from 6.4 waters in bulk to 0.2 or fewer in the bound state for both C110 routes, and from 4.2 to 0.1 for C96) and the steepest part of every profile falls where the last one or two waters leave (Figures 2J and S7A–C). The gates themselves move only transiently and only when H_2_O_2_ crosses them (Figure 2K). The H4F61 ring flips away from the bulk, the H3H113–H3′D123 latch widens and the H3′I130 plug is displaced at 3.8–4.8 Å from the cysteine (in at most 41, 65 and 46 % of the frames, respectively), and each returns to its initial state behind H_2_O_2_ (Figure S7D–F). On the top route to C110, the two replicas in which the latch opens widest at the crossing (to 5.4 Å on average, against 2.86 Å in bulk) are the two with the lowest barriers (10.3 ± 1.4 kcal/mol), while in the four replicas in which it opens only to 3.5 Å, the same route costs 15.9 ± 1.2 kcal/mol (Figure S8). The low topentry barrier is therefore conditional on a gate fluctuation and not an intrinsic property of that route.

The two cysteine sites differ in what awaits H_2_O_2_ once it arrives. At C110, the thiol has a base within reach. H3D123 approaches C110@HG to 1.60–1.74 Å in every replica for both routes, in about 13 % of the bound-state frames (Figures 2L and S7G–I). At C96 there is no equivalent. H3E97, the only carboxylate in the vicinity, never comes closer than 4.5 Å to C96@HG and sits at 8.1 ± 0.2 Å on average in the bound state, i.e. it never reaches a distance compatible with C96 deprotonation. Sulfenylation at C96 must therefore proceed either from an unassisted and strictly transient encounter complex, or after a non-negligible reorganization step bringing E97 towards the thiol group. At C110 on the other hand, a base-assisted arrangement is readily accessible as a substate of the bound state. Moreover, the target sulfur is essentially dry in the pre-reactive geometry. When an H_2_O_2_ oxygen lies within 3.5 Å of S, a water is found within 3.6 Å of S in only 0.1 ± 0.2, 3.7 ± 3.7 and 0.9 ± 1.1 % of the frames at C96, C110 top and C110 bottom entry, respectively, and a water in contact with both H_2_O_2_ and S in 0.1 ± 0.1, 1.7 ± 1.7 and 0.3 ± 0.4 % (at most 4.3 % in any replica; Figure S9). A single water can therefore reach the reactive region of C110 as a rare and transient substate, while it is never observed at C96.

In summary, reaching either cysteine costs ~12 kcal/mol, set by the desolvation of H_2_O_2_ rather than by the route, and leaves it in a nearly water-free pocket where a base is within reach at C110 (H3D123) but not at C96 (H3E97 at ~8Å). How the *S*-sulfenylation proceeds in this environment, and at what chemical cost, is addressed next.

### 2.2. *S*-sulfenylation proceeds with assistance of neighboring residues

Building on the pre-reactive H_2_O_2_–cysteine geometries identified at both C96 and C110, we next employed QM/MM simulations to characterize the reaction mechanism and energetics of the *S*-sulfenylation at each site. The environment of the cysteines in the nucleosome strongly differs from other systems on which previous theoretical studies of *S*-sulfenylation have been led [16, 17, 18], by its highly buried and hydrophobic character and the presence of only a few polar or charged residues capable of stabilizing the reaction intermediate (i.e., the thiolate moiety) within reach. Based on visual inspection of the reacting cysteines’ surroundings, three hypothetical reaction routes were investigated (Figure 3): A) direct deprotonation of the cysteine thiol by H_2_O_2_ itself, followed by a nucleophilic attack of the resulting thiolate onto the peroxonium cation (H_2_OOH^+^), forming the sulfenylated cysteine and water; B) deprotonation of the cysteine by a neighboring base, followed by attack of the thiolate onto H_2_O_2_ and release of H_2_O, with the carboxylic proton subsequently transferred to the departing oxygen, regenerating the base; C) same as in route B, but with the assistance of a nearby water molecule (tested only at C110).

**Figure 3:**
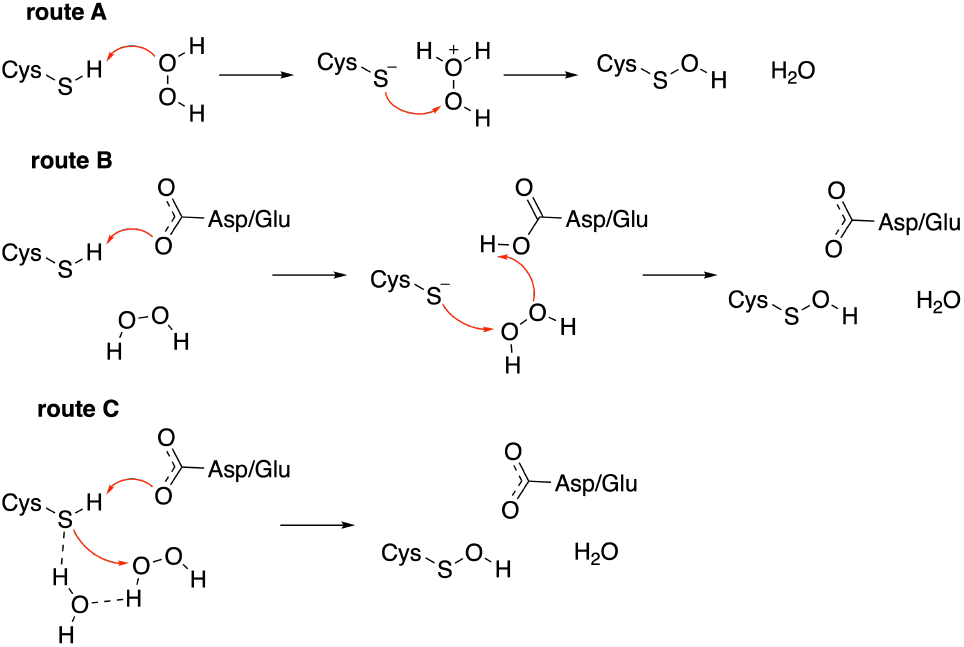
Possible mechanisms for cysteine *S*-sulfenylation in the nucleosome. Schematic of the three reaction routes tested by QM/MM, from the thiol (Cys–SH) and H_2_O_2_ to the sulfenylated cysteine (Cys–S–OH) and H_2_O. **Route A:** H_2_O_2_ is the only base. It takes the thiol proton, and the resulting thiolate attacks the peroxide. **Route B:** a neighboring carboxylate (H3E97 at C96, H3D123 at C110) takes the proton. The thiolate then attacks H_2_O_2_, and the carboxylate gets its proton back as H_2_O leaves. **Route C:** same as route B, with a water molecule relaying the proton.

#### S-sulfenylation at C96 requires the rotation of H3E97 to activate the thiol

The proximate environment of H3C96 with bound H_2_ O_2_ is mostly hydrophobic, depleted of any water molecule. H3E97 is the only closest base for available for deprotonation. In this context, only route A (deprotonation by H_2_O_2_ without assistance) and route B (deprotonation by E97) were considered for this site.

The direct activation of the thiol by H_2_O_2_ (route A) appears highly unfavorable. Indeed, no metastable state featuring the deprotonated C96 and the resulting H_2_OOH^+^ could be attained in our biased simulations, which translates into a continually growing energy profile (Figure S10A). This can be explained by the inability of H_2_OOH^+^ to be stabilized in such a hydrophobic protein environment.

The feasibility of C96 activation by H3E97 (route B) is probed here in three steps. First, E97 side chain must be rotated towards the cysteine, as it usually points away towards the loop between the H3 αN and α1 helices. This rotation costs ~7 kcal/mol, with the formation of a metastable state featuring a distance of 1.40 Å between the C96 thiol hydrogen and the closest E97 carboxylate oxygen (Figure 4A). Next, the cysteine is deprotonated by the rotated carboxylate, in a quasi-barrierless step (<1 kcal/mol; Figure 4B). Finally, the thiolate attacks the H_2_O_2_, forming the sulfenate (SO^−^) and releasing H_2_O, in a step associated with about 13 kcal/mol activation barrier (Figure 4C). The transfer of hydrogen from the reacting H_2_O_2_ oxygen to the nonreacting oxygen to form the sulfenate and water is consistent with the mechanism reported by Zeida and co-workers for the sulfenylation of methanethiolate.[16] The product formed is thermodynamically more stable than the reactant by 3 kcal/mol in this last step, suggesting that SO^−^ is more easily stabilized than the thiolate in this hydrophobic environment. A nearby phenylalanine (H4F61) seems to contribute, if only mildly, to this stabilization. Note that the final return of the carboxylic proton from the protonated E97 to the sulfenate, which converts SO^−^ into the neutral sulfenic acid and stabilize the product further, was not simulated. This step is not expected to be limiting and should not alter the overall barrier, which is set by the 13 kcal/mol peroxide attack.

**Figure 4:**
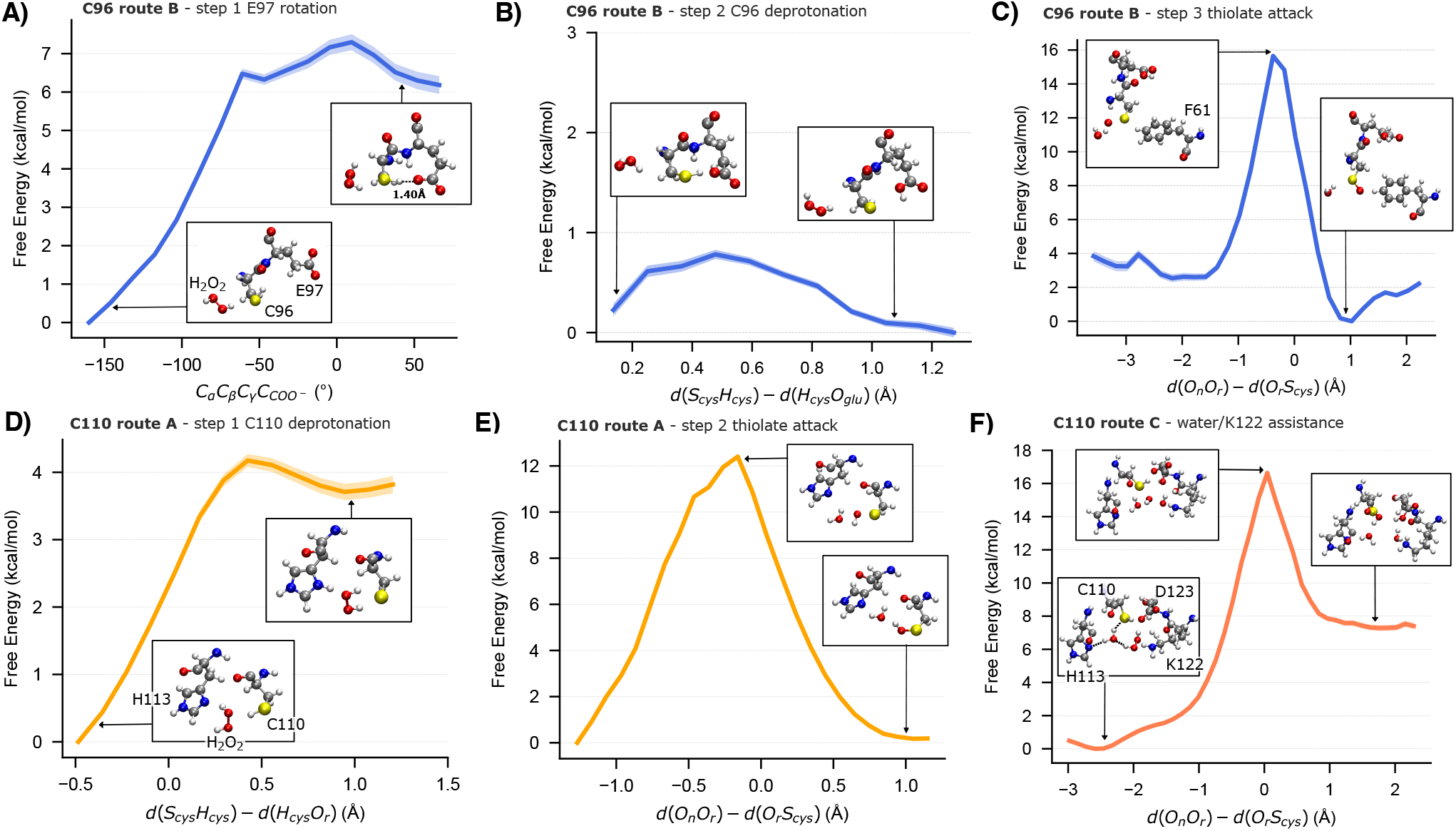
Free-energy profiles of the productive sulfenylation routes at C96 and C110. Potentials of mean force obtained by umbrella sampling along the reaction coordinate of each elementary step. Insets show representative structures of the states indicated by the arrows. **(A–C)** C96, route B three steps: **(A)** Rotation of the E97 side chain, followed along the *C*_α_*C*_β_ *C*_γ_ *C*_*COO*−_ dihedral. **(B)** Proton transfer from C96 to the E97 carboxylate, followed along *d*(*S*_*cys*_*H*_*cys*_) − *d*(*H*_*cys*_*O*_*glu*_). **(C)** Attack of the C96 thiolate on H_2_O_2_, followed along *d*(*O*_*n*_*O*_*r*_) − *d*(*O*_*r*_*S*_*cys*_), where *O*_*r*_ is the peroxide oxygen attacked by the sulfur and *O*_*n*_ the one released as water. H4F61 stabilizes the product. **(D–F)** C110. Route A two steps: **(D)** C110 is first deprotonated, the proton being transferred to H_2_O_2_ which itself transfers one of its hydrogens to H3H113, and **(E)** the resulting thiolate then attacks H_2_O_2_. **(F)** Route C: a concerted alternative in which C110 is deprotonated by H3D123, and H3K122 releases a proton for H_2_O to be released. H3H113 and a water molecule complete the hydrogen-bond network.

#### S-sulfenylation at C110 needs assistance from the H3H113-H3′D123 pair

At C110 vicinity, several polar or charged residues and rare water molecules could be involved in the reaction, notably the H3H113-H3′D123 pair that can be released upon H_2_O_2_ entry as described above. The three mechanistic routes have thus been investigated here.

The deprotonation of C110 by H_2_O_2_ itself (route A) occurs with a free energy barrier of ~4 kcal/mol (Figure 4D) but the proton is actually immediately shuttled to the nearby imidazole nitrogen of H3H113, leading to an intermediate featuring the thiolate, the neutral H_2_O_2_, and the protonated H113. The thiolate attacks the H_2_O_2_, proceeding with a free energy barrier of ~13 kcal/mol to form the sulfenic acid in a kinetically limiting step (Figure 4E). The hydroxide (OH^−^) released from H_2_O_2_ upon sulfenic acid formation finally captures back the proton previously transferred to H3H113, releasing H_2_O as a by-product and regenerating the neutral H113.

The deprotonation of C110 by the nearby aspartate (route B, H3′D123) is quasi barrierless (~3 kcal/mol), but the resulting cysteinate intermediate is not stable and instantaneously reverts, owing to its poor stabilization (Figure S10B). In addition, a strong hydrogen bond between the H3H113 imidazole nitrogen and the peroxide proton likely increases the nucleophilicity of H_2_O_2_ oxygen atom, which is positioned close to the cysteinate (S–O distance of ~2.19 Å), further disfavoring thiol oxidation along this route at C110.

For route C, a water molecule entering the reactive region forms a hydrogen bond with the peroxide proton (O_wat_ – H_perox_ distance of ~1.70 Å; Figure 4F). This water molecule simultaneously forms hydrogen bonds with the H3H113 imidazole nitrogen (H_wat_ –N_his_ distance of ~1.90 Å) and the C110 thiol sulfur (H_wat_ –S_cys_ distance of ~2.57 Å), forming a hydrogen-bond network that assists deprotonation of the cysteine by the nearby aspartate. Sulfenylation along this route proceeds with an overall free energy barrier of ~17 kcal/mol. Upon sulfenylation, the hydroxide anion released from the peroxide abstracts a proton from a nearby lysine (H3K122), forming H_2_O as a by-product. This leaves H3K122 neutral, and it must therefore be re-protonated, most likely from the bulk solvent through a water-mediated proton relay. This step was not simulated. It occurs after the ratelimiting step and should be thermodynamically favorable, since a neutral lysine is a strong base, so we do not expect it to add to the overall barrier.

Overall, these results show that *S*-sulfenylation can proceed at both cysteines without bulk water. At C96, the only viable route is activation by H3E97, which must first rotate towards the thiol (~7 kcal/mol) before deprotonating it and allowing the thiolate to attack H_2_O_2_ (12 kcal/mol), whereas direct activation by H_2_O_2_ fails because H_2_OOH^+^ cannot be stabilized in this hydrophobic environment. At C110, the base is already in place: the reaction through both H3H113 (route A, ~17 kcal/mol) and through H3′D123 with assistance of a water molecule (route C, ~17 kcal/mol) seems plausible, however, the latter is more favourable given the high likelihood of the reversibility of step 1 of the former. The deprotonation by H3′D123 alone (route B) on the other hand fails owing to the instability of the cysteinate intermediate without water. The chemical barriers are therefore of the same order at the two sites, and what differs is the cost of the reorganization needed to reach the reactive arrangement. We note that each step was described by one-dimensional umbrella sampling along a single reaction coordinate, so other degrees of freedom that may contribute to the reaction (e.g., the orientation of the neighboring residues, the position of the water molecule, or the concerted motion of several proton transfers) were not biased. The transition states identified here may therefore not be the lowest-energy ones, and the barriers should be regarded as upper estimates. These profiles nevertheless provide a first, necessarily simplified, view of the reaction and identify the residues likely to take part in it (H3E97 at C96; H3H113, H3′D123, H3K122 and a water molecule at C110).

### 2.3. C96 and C110 oxidation perturb distinctly the nucleosome core particle dynamics

After describing the formation of *S*-sulfenylation at C96 and C110, we next investigated how this oxidative modification impacts nucleosome dynamics and histone–DNA contacts. The systems with sulfenylated C96 and C110 are noted H3.1soh and H3.2soh, respectively. For C110, we further compared these effects to *S*-nitrosylation (H3.2sno simulated here, given its role as a reaction intermediate toward *S*-glutathionylation) and to *S*-sulfonylation (H3.2ocs), whose destabilizing structural effects were previously described [14]. Because the timescales accessible to all-atom MD simulations cannot capture the entire conformational landscape of the full-length systems, each system was additionally simulated with truncated histone tails. This approach strongly simplifies the model to overcome the sampling limitations imposed by the intrinsically disordered nature of the histone tails, thereby improving the sampling of slow motions in the nucleosome architecture. Comparing full-length and tailless systems also reveals to what extent the effects of the modifications depend on the histone tails.

#### Cysteine oxidation perturbs DA dynamics in a siteN-, chemistry- and tail-dependent manner

To assess how oxidation of the H3 cysteines impacts nucleosomal DNA dynamics, we first monitored DNA end-to-dyad distances to probe DNA unwrapping events along the MD simulations (Figure 5A–C).

**Figure 5:**
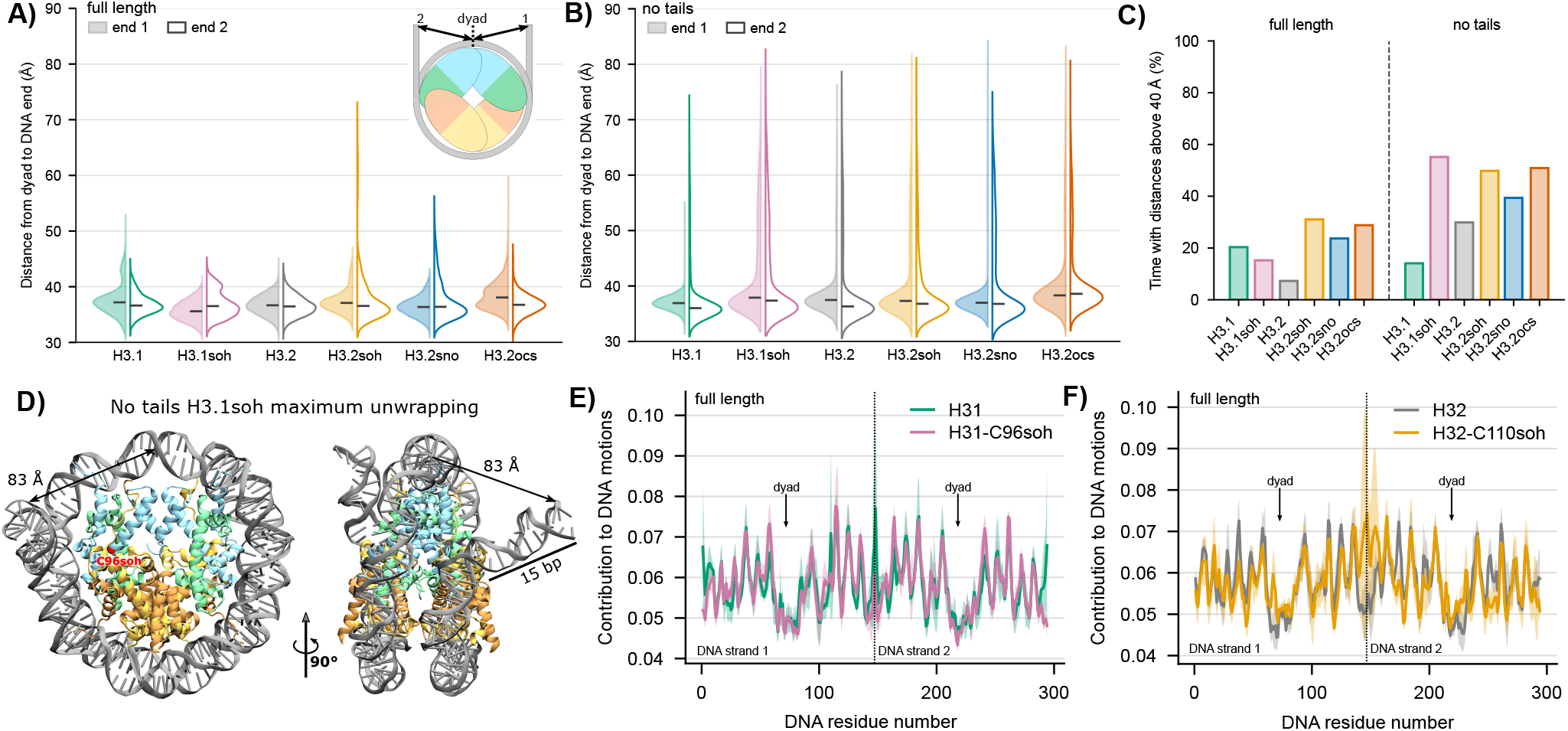
DNA unwrapping and dynamics of nucleosome core particles bearing cysteine post-translational modifications. (A) Distribution of end-to-dyad distances during MD simulations of full-length nucleosome core particles containing H3.1 (green), H3.1 bearing *S*-sulfenylated C96 (pink), H3.2 (grey), H3.2 bearing *S*-sulfenylated C110 (yellow), H3.2 bearing *S*-nitrosylated C110 (blue), or H3.2 bearing *S*-sulfonylated C110 (orange). The inset illustrates the monitored distances. (B) As in (A), for nucleosome core particles with truncated histone tails. (C) Percentage of simulation time during which at least one end-to-dyad distance exceeds 40 Å. (D) Conformation corresponding to the maximum unwrapped state sampled for the nucleosome core particle containing *S*-sulfenylated H3C96, shown in face (left) and side (right) views; the modification site is highlighted in red. (E) PCA-based per-residue contributions to DNA collective motions during MD simulations of full-length nucleosome core particles containing canonical or *S*-sulfenylated H3.1. (F) As in (E), for canonical or *S*-sulfenylated H3.2.

In the full-length nucleosome, both DNA ends remain closely wrapped around the histone core for the unmodified systems and for *S*-sulfenylated H3.1 (H3.1soh), with only slight breathing motions and dyad-to-end distances rarely exceeding 40 Å (Figure 5A). In contrast, all three C110-modified H3.2 systems (H3.2soh, H3.2sno, and H3.2ocs) sample much larger dyad-to-end distances, reflecting more pronounced DNA breathing and a few unwrapped conformations. This trend is well-illustrated by the fraction of simulation time spent with a dyad-to-end distance above 40 Å. The three C110-modified systems all showed markedly higher values than the unmodified H3.2 (20–30% vs. 8% for the control), while H3.1soh remained close to its own unmodified counterpart (18–20%; Figure 5C). Of note, the enhanced breathing is more pronounced on the side of the modification site (dyad-to-end2 distance), except for *S*-sulfonylation.

Removing the histone tails increases the sampling of unwrapping events in all six systems, which then reach comparably large maximal distances (Figure 5B). In this tailless context, H3.1soh shows the highest percentage of time above the 40 Å threshold across the whole dataset, exceeding even the C110-modified systems (Figure 5C). The maximum unwrapping observed reaches an end-to-dyad distance of ~80–85 Å, that translates into an opening of the DNA up to 15 DNA base pairs (SHL7 to SHL5.5) out of the nucleosome wheel plane (Figure 5D). DNA unwrapping is a slow event, and 5 replicas of 2 µs per system are likely insufficient to fully sample it in the full-length systems, where the flexibility of the disordered tails slows convergence. However, sampling alone may not account for the difference between the full-length and tail-truncated trends, as the histone tails might also directly modulate how the effect of the modifications propagates to the DNA, a point we address in the following sections.

PCA-based per-residue contributions to DNA collective motions show that, while no significant difference is observed between *S*-sulfenylated C96 and its canonical reference, sulfenylation of C110 destabilizes the DNA around the dyad (Figure 5E–F), as we previously described for *S*-sulfonylation [14]. The dyad is well-described as the most stable region of nucleosomal DNA, and its destabilization has been linked to nucleosome disassembly processes [19]. Interestingly, this effect is not observed upon *S*-nitrosylation, suggesting that the specific chemistry of the modification can play a role in the structural perturbation, as *S*-sulfenylation and *S*-sulfonylation both introduce an oxygenbased modification on the sulfur atom while *S*-nitrosylation exhibits a distinct nitroso (S–N) moiety. In addition, although neither cysteine directly contacts the DNA, C110 lies closer to the dyad (~15 Å below) than C96. Hence, they might act on distinct regions of the nucleosome and exhibit different structural signatures.

As expected from the different trends observed for DNA unwrapping, removing the histone tails drastically changes the DNA flexibility profile of the H3.1 variant, with much higher contributions of the DNA ends than for its fulllength counterpart (Figure S11A). Without the tails, C96 *S*-sulfenylation increases DNA motions at SHL ±0.5, which is compensated by a stiffening at other SHLs except the DNA ends. These enhanced dyad motions might arise from the extensive DNA unwrapping observed in this system. Indeed, the N-terminal end of the H3 αN helix (R40–A47, also known as the H3 latch [20]) is sandwiched between the two DNA gyres, simultaneously contacting the DNA ends and the DNA at SHL ±0.5. The release of the DNA ends perturbs this bridge, which could propagate the destabilization towards the dyad. In contrast, DNA motions for systems with H3.2 show a lower increase at these SHL compared to the full-length trends (Figure S11B).

Overall, H3 cysteine oxidation perturbs nucleosomal DNA dynamics in a site-, chemistry- and tail-dependent manner. The effects of C96 *S*-sulfenylation on both the DNA ends and the dyad emerge only in the absence of the histone tails, while C110 modifications favour DNA unwrapping and, for oxygen-based modifications, destabilization of the dyad even in the full-length nucleosome.

#### C96 and C110 oxidation redistribute key contacts at histone–histone and DNA–histone interfaces

Having shown that C96 and C110 oxidation affect DNA dynamics differently, we next assessed their effects on histone–histone and histone–DNA contacts by probind interactions networks with InterMap [21]. We specifically focused on the H3–H3′ interface (which harbors C110 and has been linked to nucleosome disassembly), the H3–H4 interface (which harbors C96), and the (H3–H4)_2_–H2A/H2B and (H3–H4)_2_–H2A′/H2B′ interfaces (also linked to disassembly [22]) (Figure 6A).

**Figure 6:**
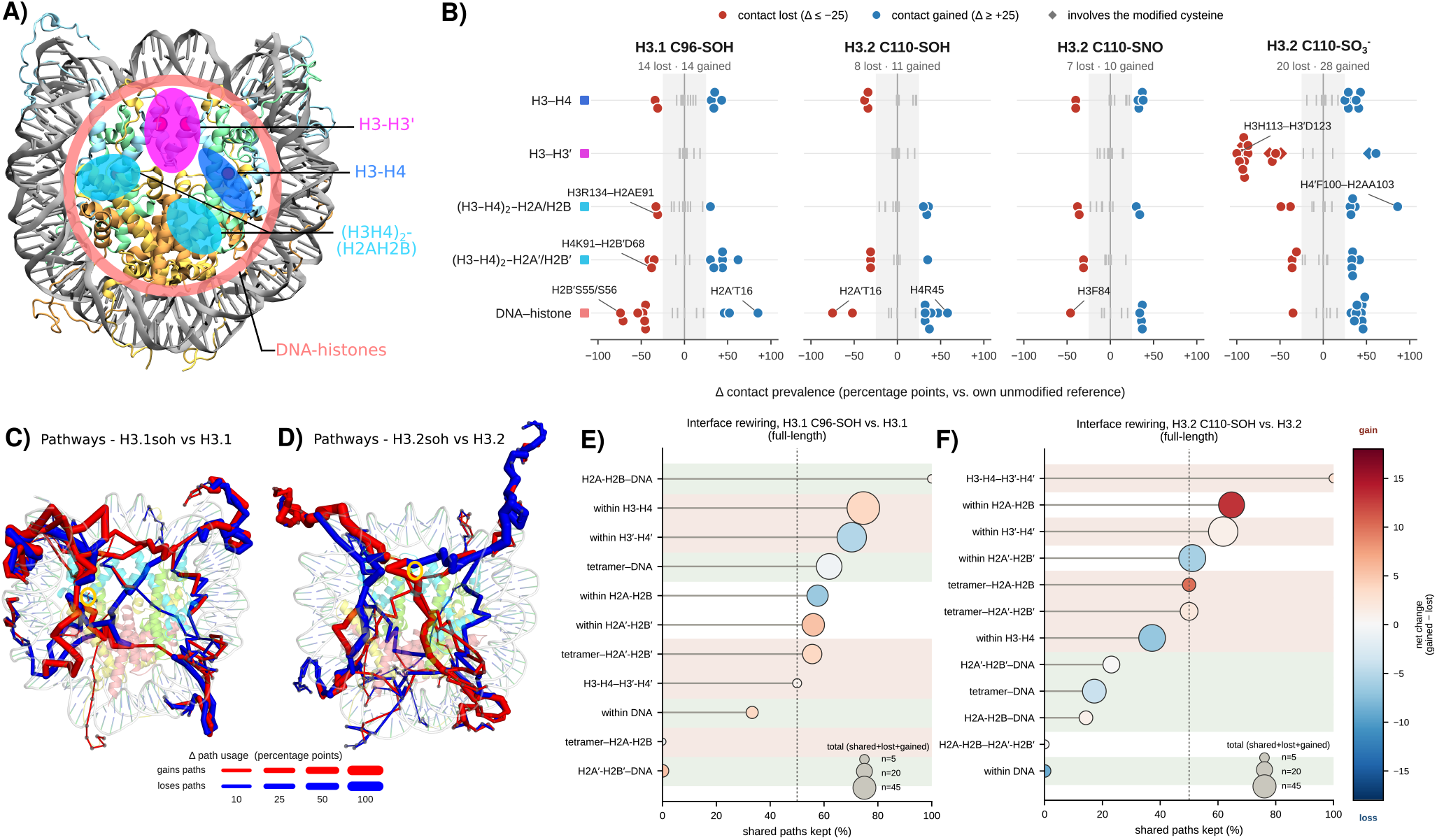
PTM-induced remodeling of intra-nucleosome contacts and communication pathways. (A) Structural overview of the nucleosome core particle with the interfaces analyzed in (B) mapped onto the structure: H3–H3′ (magenta), H3–H4 (dark blue), (H3–H4)_2_–H2A/H2B tetramer–dimer (cyan), and DNA–histone (salmon). (B) Change in intermolecular contact prevalence (Δ, percentage points of MD frames in close contact) upon modification, each system being compared to its own unmodified reference: H3.1 for C96 *S*-sulfenylation, H3.2 for C110 *S*-sulfenylation, *S*-nitrosylation, and *S*-sulfonylation. Each point is one residue pair, and each row one interface (colored squares as in A). Contacts lost (Δ ≤ −25) are shown in red, contacts gained (Δ ≥ +25) in blue, and smaller changes as grey ticks within the shaded band. Diamonds show contacts directly involving the modified cysteine. The number of contacts lost and gained beyond the threshold is given under each system name, and the contacts discussed in the text are labeled. For each interface, the 15–19 most perturbed residue pairs are shown, restricted to the histone-fold core with N-/C-terminal tails excluded. Individual values are given in Figure S12. (C, D) Difference maps of the ComPASS optimal-communication network for full-length H3.1 C96-SOH versus H3.1 (C) and H3.2 C110-SOH versus H3.2 (D). Each stick is an edge of the top-10% shortest-paths network, connecting consecutive residues along a path. Red edges are used by a larger fraction of top paths in the modified system than in its reference, and blue edges by a smaller fraction. Stick width scales with the change in path usage. (E, F) Interface rewiring for H3.1 C96-SOH (E) and H3.2 C110-SOH (F). For each pair of nucleosome regions, the bubble position gives the percentage of reference paths kept in the modified network, its color the net change in path number (gained minus lost), and its size the total number of paths involved (shared + lost + gained). Shaded rows distinguish histone–histone (pink) from DNA-related (green) region pairs.

At the H3–H4 interface, *S*-sulfenylation of C96 rewires the local contact network (Figure 6B and S12B). Several contacts near the modification site are reinforced (e.g. H3E73–H4D24, H3P66–H4N25, H3R63–H4T30), while others are weakened, including the H3D81–H4K79 (−31%) and H3R63–H4R36 (−34%). The direct C96–H4 contacts themselves (H3C96–H4L58/F61/L62) are unchanged, suggesting that *S*-sulfenylation perturbs the surrounding H3– H4 dimer packing without affecting the cysteine’s nearby hydrophobic environment. While the more distant H3–H3′ interface remains unaffected, the local rearrangement around C96 propagates to other more distal interfaces (Figure 6B and S12C–E). Indeed, at the (H3–H4)_2_–H2A/H2B and (H3– H4)_2_–H2A′/H2B′ interfaces, C96 *S*-sulfenylation disrupts several important salt bridges and hydrogen bonds. The H3R134–H2AE91 salt bridge is weakened (−31%), releasing H2A E91, which belongs to the H2A/H2B acidic patch. In addition, the H4K91–H2B′D68 salt bridge (−38%), which bridges the two halves of the nucleosome, is consistently weakened in all five replicas. The loss of the H4G42– H2A′L115 contact (−35%) reflects the release of the H2A′ C-terminal tail (residues 110–117) from its position near the dyad, where it usually packs against the H3 αN (L48, I51) and α3 (N108, I112, V117) helices and H4 K44. Monitoring the number of native contacts between the H2A C-terminal tail and the (H3–H4)_2_ tetramer illustrates this release, which is persistently observed in one H3.1soh replica and only transiently in the H3.2soh and H3.2sno systems (Figures S13– S15). Importantly, these changes do not translate into a breathing of the H2A/H2B dimers, as the contacts made by their structured cores with the tetramer (mostly in the H4– H2B four-helix bundle) remain stable in all replicas of all six systems. C96 oxidation therefore rewires specific native contacts well beyond its own interface, including key salt bridges, and propagates to the H2A/H2B dimers on both sides of the tetramer, without dramatically loosening the core of the dimer–tetramer interface. The most pronounced long-range effect is, however, seen at the DNA–histone interface (Figure 6B and S12F). C96 *S*-sulfenylation weakens several arginine/lysine–DNA contacts, including H4′R36 at SHL-1.5 (−54%), H2AR29 at SHL-4.5 (−48%), H2A′H31 at SHL+4.5 (−46%), and also notably H4R45 (−45%) at SHL-0.5, an arginine anchor directly adjacent to the dyad. Most of these residues nevertheless remain bound to the DNA and rather shift their contacts to neighbouring nucleotides (e.g. H4R45 remains engaged with the DNA 100% of the time). The clearest losses are those of H2B′S55 and H2B′S56 at SHL+5.5 (−71% and −74%). Some DNA contacts are reinforced elsewhere (e.g. H2A′T16 at SHL+5, H3′R72 at SHL-2), again suggesting a redistribution of histone–DNA contacts rather than to a global loss.

The trends are markedly different for C110. At its own interface (H3–H3′), *S*-sulfenylation and *S*-nitrosylation leave the contacts unchanged, whereas *S*-sulfonylation (H3.2ocs) destabilizes the entire four-helix bundle interaction network with a complete loss of the H3H113–H3′D123 key contact (Figures 6B and S12C), consistent with our previous report [14]. At the more distal (H3–H4)_2_–H2A/H2B interface, all three C110 modifications show a loss of the H3R134–H2AE91 salt bridge, thus releasing the acidicpatch residue H2AE91, and the H2AR99 contacts with H4′G94 and H4′R95 (Figure 6D). The opposite (H3–H4)_2_– H2A′/H2B′ interface remains unperturbed by sulfenylation and nitrosylation, and is only affected by sulfonylation, with loss of H3–H2A′ contacts and gain of several H4– H2A′/H2B′ contacts. This points to an asymmetric propagation of the C110-induced perturbation across the two sides of the nucleosome. At the DNA–histone interface the effect of C110 oxidation contrasts strikingly with that of C96 sulfenylation, with the H4R45 contact at SHL-1 being reinforced by all three C110 modifications, as is the H2AR29 contact at SHL-4 (Figure 6F). C110 oxidation thus acts primarily on the DNA and on one side of the histone core, without the extensive rewiring of histone–histone contacts seen for C96. The dyad DNA is perturbed even though its main histone anchors are not lost.

In the tailless particles, the histone–DNA contacts perturbed by C96 *S*-sulfenylation are located at the DNA ends (H3′R49 at SHL −6.5, −39% and H3′T45 at SHL −7, −45%) instead of being spread along the whole sequence, whereas the H3– H3′ interface remains as unaffected as in the full-length system (Figure S16). These end contacts include the outermost arginine anchors (H3 R49 of the αN helix, SHL ±6.5), which are released from the DNA 33–56% of the time in all modified tailless systems, against 15–17% for unmodified H3.1 and H3.2, while their release remains below 14% in all full-length systems regardless of the modification. In the full-length nucleosome, these anchors are flanked by the H3 N-terminal tail, whose K36 and K37 residues directly contact the DNA ends at SHL ±7, a restraint that is lost upon truncation. The different unwrapping trends observed between full-length and tailless systems therefore mainly stem from the removal of these tail-mediated restraints, which renders the outermost anchors sensitive to the modifications, an effect buffered by the tails in the full-length nucleosome. This also suggests that the long-range action of C96 *S*-sulfenylation on histone–DNA contacts works cooperatively with the histone tails, more than for the C110 modifications, whose tailless signature mostly reflects their enhanced DNA unwrapping.

Together, these results show that C96 *S*-sulfenylation specifically weakens key inter–histone salt bridges (H4K91– H2B′D68, which holds the two halves of the nucleosome together, and H3R134–H2AE91 at the acidic patch) and redistributes histone–DNA anchoring contacts in a different manner than C110 oxidation. These changes occur without any apparent breathing of the H2A/H2B dimer cores. The only large-scale histone event observed, that is, the release of the H2A′ C-terminal tail from the dyad region, is rare but present. This supports the idea that distinct destabilization mechanisms are at play for the two sites. Since most of these changes arise far from the modified cysteines, they likely reflect allosteric effects propagated from the modification sites rather than solely local consequences. The comparison with the tailless systems further reveals cooperativity between C96 *S*-sulfenylation and the histone tails. Once the tails are removed, the long-range action of the modification on histone–DNA contacts is abolished, and the loss of the tail restraints at the DNA ends exposes the outermost arginine anchors, whose release becomes the dominant signature of this modification.

#### Both modifications rewire communication through the histone tails and the H3/H3′ latch

To investigate how the perturbations induced by cysteine oxidation propagate from the modification sites to distal interfaces, we characterized the long-range communication networks in the different systems using ComPASS [23].

Comparing the top pathways connecting distant regions of the nucleosome for each modified network with its unmodified reference, we found that both modifications extensively rewire the communication networks (Figures 6C– F and S17). The C96 *S*-sulfenylated network retains only 46.5% of the reference network, while the C110-modified networks even less (28.0% for *S*-sulfenylation, 40.7% for *S*-nitrosylation). Notably, neither cysteine lies on any top pathway in any system, suggesting that the two sites are not communication hubs themselves but reorganize the traffic around them.

The two sites act on different parts of the network (Figure 6C–F). C96 *S*-sulfenylation mainly changes the communication between histones, at interfaces whose loosening is required for nucleosome disassembly. The pathways linking the (H3–H4)_2_ tetramer to one H2A/H2B dimer are almost entirely rewired, half of those linking the two H3–H4 halves of the tetramer are lost, and the direct link between H4R45 and the dyad DNA disappears. This is in line with the weakening of the H4K91–H2B′D68 and H3R134–H2AE91 salt bridges and with the redistribution of the H4R45–DNA contacts described above. C110 *S*-sulfenylation shows the opposite pattern. The histone–DNA pathways are the most impacted, with all four DNA interfaces keeping less than 25% of their pathways, while the H2A/H2B and H3′–H4′ dimers keep more than 60%, consistent with the destabilization of the DNA around the dyad observed in the PCA. *S*-nitrosylation at the same site is milder, as it retains more of the reference network (40.7% against 28.0%) and does not change the number of top pathways (Figure S17).

Two common features stand out from comparison with the tailless simulations. First, the histone tails are used as shortcuts. In the unmodified systems, top pathways are of comparable length in the full-length and tailless versions, whereas upon *S*-sulfenylation the full-length pathways become markedly shorter than their tailless counterparts at both sites (Figure S17). The number of top pathways drops accordingly, from 1424 to 844 for C96 *S*-sulfenylation and from 1103 to 742 for C110 *S*-sulfenylation. This tailmediated wiring is consistent with the cooperativity between C96 *S*-sulfenylation and the histone tails observed for the histone–DNA contacts. Secondly, the H3/H3′ latch, which bridges the DNA ends to the dyad-proximal DNA and gates DNA unwrapping [20], becomes an almost obligatory hub upon modification at both sites. It wires 43.5% of the pathways in unmodified H3.1 against 80.4% after C96 *S*-sulfenylation, and 6.7% in unmodified H3.2 against 100% after C110 *S*-sulfenylation and *S*-nitrosylation (Figure S17).

Overall, neither C96 nor C110 oxidation places the modified cysteine on the communication network, but both rewire the pathways through the histone tails and the H3/H3′ latch. For C96 *S*-sulfenylation, this rerouting targets the histone–histone interfaces, in line with the weakened dimer– tetramer salt bridges and the tail-dependent redistribution of histone–DNA contacts, whereas for C110 oxidation it targets the histone–DNA contacts, in line with the destabilization of the DNA at the dyad.

## 3. Discussion

Redox signaling has emerged as a way to regulate DNA compaction, with oxidation of the buried H3 cysteines acting as a molecular switch for chromatin remodeling and for cancer-associated histone exchange and chemoresistance [6, 11]. Yet how reactive species reach these shielded thiols, how the chemistry proceeds in a hydrophobic environment, and how the modification reshapes the nucleosome dynamics remain elusive from a structural point of view.

Our simulations show that neither cysteine is reachable by diffusion alone. Reaching either site costs 9 to 18 kcal/mol depending on the replica, and 12–14 kcal/mol on average, without any significant difference between C96 and the two entry pathways to C110. This cost is mostly paid in desolvation, as H_2_O_2_ must shed its entire first hydration shell to reach the cysteines while the gates lining the pathways (H4F61 at C96 and H3H113-H3′D123 or H3′I130 at C110) open transiently as H_2_O_2_ crosses them and only tune the cost from one replica to the next. Rather than a limitation, this cost might be functionally beneficial since an easily accessible, low-barrier route would leave both cysteines constantly exposed to oxidation, favoring over-oxidation towards an irreversible *S*-sulfonylation rather than a signaling-compatible sulfenylated state. All access pathways pass through a pocket acting as an antechamber, where the elongated geometry of H_2_O_2_ allows it to interact simultaneously with vicinal backbone carbonyl oxygens and with the cysteine itself. Water cannot achieve this, and the resulting selectivity may itself contribute to the oxidant-sensing specificity of these sites. Notably, the residues of both pockets are essentially H3 and H4 residues, which suggests that this access mechanism could already operate in isolated (H3–H4)_2_ tetramers or H3– H4 dimers. Interestingly, the top entrance to C110, gated by H3H113 and H3′D123, lies at the H3–H3′ interface where C110, H3H113 and the axial H3L126 form the Cu(II) binding site of the nucleosome copper reductase [7, 24], suggesting that this cryptic pocket may also serve as an access route for metal ions [25].

The chemistry is constrained by the same burial. At both sites, *S*-sulfenylation requires a neighboring residue to deprotonate the cysteine, since neither is accessible to bulk solvent for a solvent-mediated process: H3E97 at C96, and either H3H113 or H3D123 at C110. A single wellpositioned residue therefore substitutes for the catalytic role otherwise played by the aqueous environment. Remarkably, the resulting barriers (~17–18.5 kcal/mol) are of the same order as those reported for cysteine oxidation by H_2_O_2_ in solvent-exposed or highly polar environments [16, 17, 18], although our cysteines react with no bulk water available to stabilize the reaction intermediates. This remain substantial, in line with the idea that *S*-sulfenylation of H3C96 and H3C110 should not be an easy event, as an easily crossed barrier would leave both cysteines constantly reactive towards H_2_O_2_, favoring over-oxidation to irreversible *S*-sulfonylation rather than a signaling-compatible sulfenic acid state. The reaction free energies we obtain are close to thermoneutral or unfavorable (~0 kcal/mol for C96 route B, +4 and +8 kcal/mol for C110 routes A and C). This likely reflects that the product states reached here are not the true final states, as several relaxation steps were not simulated. The released water or hydroxide are still in the pocket (for all routes), H3K122 is left deprotonated instead of being re-exposed to the solvent and protonated back (route C at C110), and the carboxylic proton of H3E97 is not returned to the sulfenate (route B at C96). These steps occur after the rate-limiting step and are expected to stabilize the products, so the barriers are not affected, but the reaction free energies reported here should be regarded as upper estimates.

No direct measurement of the reaction rate at H3C96 or H3C110 is available, but our barriers agree qualitatively with the slow reaction of non-catalytic protein thiols with H_2_O_2_, several orders of magnitude below the specialized peroxiredoxins [26, 27]. Oxidation of such modestly reactive thiols has been proposed to occur indirectly in cells, through redox relay via an already-oxidized peroxiredoxin or thioredoxin [27]. This would however require an interface between the target cysteine and the relay protein, which the same burial that imposes a dedicated H_2_O_2_ access pathway precludes here. Direct reaction with H_2_O_2_ therefore appears to be the only mechanistically accessible route at these two sites, even if it is intrinsically slow. We finally note that these mechanisms were described with one-dimensional umbrella sampling, which cannot probe concerted events but provides a first picture of the reaction and identifies the residues important for it to proceed in such a hydrophobic environment.

Our results also provide a structural rationale for the role of H3.1 C96, whose *S*-sulfenylation was shown to promote H3.1 eviction and replacement by H3.3 in cancer cells [6]. In our simulations, C96 *S*-sulfenylation does not trigger nucleosome disassembly by itself. Indeed, the DNA ends remain wrapped in the full-length particle, and the structured cores of the H2A/H2B dimers stay firmly docked onto the tetramer. Rather, the modification acts on elements that are key for the binding and action of chromatin remodellers and chaperones. It releases the acidic-patch residue H2AE91 from its salt bridge with H3R134, weakens the H4K91– H2B′D68 salt bridge that couples the two halves of the nucleosome, and is accompanied by a transient release of the H2A C-terminal tail at the dyad. Both the acidic patch and the H2A C-terminal region are recognition elements for chromatin remodelers, and truncating the latter is known to destabilize the nucleosome and accelerate histone exchange [28, 29, 30, 31]. This C-terminal tail release was however observed in only one of the five replicas of the H3.1soh system, and coarse-grained simulations or dedicated experimental measurements would be needed to build reliable statistics and validate this hypothesis. Such a priming effect would be particularly relevant for H3 variant exchange, which requires the full disassembly of the nucleosome and therefore the action of remodellers and chaperones. The effect of C96 *S*-sulfenylation on the DNA is mediated by the histone tails, suggesting that the redox signal is propagated to the DNA through them rather than reaching it directly. It is pertinent to note that, *S*-sulfenylation and *S*-nitrosylation of C110 do not reproduce these trends, consistent with the experimental observation that the C96S mutation, or the use of H3.2 instead of H3.1, abolishes the effect of H_2_O_2_ on H3.3 incorporation [6]. C110 *S*-sulfenylation acts instead on DNA unwrapping and the dyad, and likely leads to a different biological outcome, perhaps closer to the chromatin opening reported for its *S*-glutathionylation [12, 13]. This distinction is also visible in the communication networks, as the pathways rewired by C96 *S*-sulfenylation mainly link the tetramer to the H2A/H2B dimers, in line with the weakened histone–histone contacts, whereas those rewired by C110 oxidation mainly involve the DNA, even though C96 *S*-sulfenylation weakens more individual histone–DNA contacts. This suggests that C110 modifies the dynamic coupling of the DNA to the histone core rather than specific histone–DNA contacts. *S*-nitrosylation, finally, has milder effects throughout, in line with the hypothesis that it could act as a simple intermediate towards *S*-glutathionylation.

Altogether, our results describe how histone H3 is oxidized within the nucleosome and explain, at the structural level, its role as a redox switch. Reaching the buried cysteines requires defined access pathways and a significant energy cost, mostly set by the desolvation of the oxidant, and the chemistry itself needs the assistance of well-placed residues within an otherwise hydrophobic pocket, readily available at C110 (H113-D123′) but only after a reorganization step at C96 (E97). Oxidation is therefore slow and controlled, which might help preserve the reversible sulfenic acid state and prevent irreversible over-oxidation. Once formed, the modification is felt well beyond the reaction site and rewires contacts and communication networks in a site-specific way. C96 *S*-sulfenylation primes the nucleosome for remodelling without destabilizing the octamer core, whereas C110 oxidation acts on the DNA around the dyad. Since cancer cells commonly sustain elevated levels of reactive oxygen species as a consequence of their metabolic rewiring, such redox-sensitive histone switches could link the metabolic state of the cell to chromatin structure and epigenetic modulation. By resolving, at the atomic level, how an oxidant reaches a buried histone residue, how the chemistry proceeds there, and how the resulting modification propagates through the nucleosome, this work provides a framework to investigate other oxidative histone modifications and to guide their experimental characterization.

## 4. Methods

All MD simulations were performed with the AM-BER20 suite of programs, interfaced with TeraChem 1.97B for the QM/MM. VMD [32] and PyMOL [33] were used for visualization and rendering of figures. MD ensembles of full-length unmodified and *S*-sulfonylated H3.2 nucleosomes (5×2 µs each) were taken from a previous work [14]. The total sampling amounts to ~55 µs for the study of H_2_O_2_ access (unbiased MD with free H_2_O_2_, relaxation runs after steered MD, and umbrella sampling), ~100 µs for the modified and unmodified (control) nucleosomes (in addition to the 20 µs taken from [14]), and ~1 ns for the QM/MM relaxation and umbrella sampling.

### 4.1. Systems setup

Starting systems were built from the crystal structure of the nucleosome core particle featuring an α-satellite DNA sequence and full histone tails (PDB ID 1KX5 [34]). The three missing residues (PEP) at the N-terminus of both H2B copies were reconstructed using VMD and the LEaP module of AMBER20. Systems with truncated tails were taken from ref. [20], while systems with H3.1 were generated by *in silico* mutation of H3.2S96 to C96. The AMBER ff14SB force field [35] was used in combination with bsc1 [36], the CUFIX corrections for non-bonded terms [37], and the TIP3P water model [38].

#### Preparation of the systems with H_2_**O**_2_

Using the packmol software [39], 200 H_2_O_2_ molecules were randomly placed around the H3.1 system at a minimum distance of 10 Å between molecules. The system was placed in a water box with a truncated octahedron shape,a 15 Å buffer, and a 0.15 M NaCl concentration. Force field parameters for H_2_O_2_ were generated following the antechamber protocol. The resulting systems consisted of ~310,000 atoms.

#### Preparation of the oxidized nucleosomes

Oxidative modifications were introduced on the first copy of histone H3 using the LEaP module of AMBER20. Three systems were generated with and without histone tails: (i) *S*-sulfenylated H3.1C96, (ii) *S*-sulfenylated H3.2C110, and (iii) *S*-nitrosylated H3.2C110. Tail-less systems, either unmodified or bearing *S*-sulfonylated H3C110, were also generated to enable comparison with their full-length counterparts, for which data was obtained from a previous study [14]. Parameters for the *S*-sulfenylated cysteine were generated in-house as described for *S*-sulfonylation in previous work [14], and those of the *S*-nitrosylated cysteine were taken from the literature [40]. Each modified system was placed in a water box with a truncated octahedron shape, a 20 Å buffer, and a 0.15 M NaCl concentration. The resulting systems consisted of ~400,000 and ~150,000 atoms for the systems with and without histone tails, respectively.

### 4.2. Molecular dynamics (MD) simulations

The starting structures were first optimized in 4 steps with decreasing position restraints on the nucleosome core particle atoms, from 20 kcal/mol/Å^2^ to 5 kcal/mol/Å^2^. In each of these 4 steps, a minimization run of 10,000 steps (5,000 steps of steepest descent followed by 5,000 steps of conjugate gradient) was followed by a short equilibration run of 20 ps at 100 K. This procedure allowed the structure to be smoothly optimized within its environment. A final 10,000-step minimization without restraint was then followed by a 20 ps thermalization to increase the temperature to 300 K, and a 500 ps equilibration in the NVT ensemble to equilibrate the solvent. The system was then relaxed in NPT for 100 ns. A 2 µs production run was then performed using a 4 fs timestep, allowed by the use of the HMR approach on the solute atoms [41, 42]. Bonds involving hydrogen were constrained using the SHAKE algorithm [43]. Temperature and pressure were kept constant (300 K, 1 bar) using the Langevin thermostat with a 2 ps^−1^ collision frequency and the Berendsen barostat with isotropic position scaling and a pressure relaxation time of 1 ps. An 8 Å cutoff was used for non-bonded terms, and long-range electrostatics were treated using the Particle Mesh Ewald approach [44]. For each system, five replicas were produced with random starting velocities, resulting in a total of 10 µs per system. RMSD of the histone core and of DNA was monitored along all simulations to check convergence (Figures S18 and S19).

### 4.3. Steered MD

Steered MD simulations were performed to generate conformations featuring H_2_O_2_ in close proximity to H3C96 or H3C110, which was further used for QM/MM simulations. Initial conformations were chosen from unbiased MD simulations featuring H_2_O_2_ at its minimum distance to the target cysteine. A pulling force constant of 7 kcal/mol/Å^2^ was then applied to reduce the distance between H_2_O_2_@O1 and the cysteine @SG from 5 Å to 2.5 Å, along a 5 ns run. The final conformation was then submitted to a 1 µs unbiased MD run, performed in 10 replicas, to evaluate the residence time of the molecule near each cysteine and its possible exit pathways. The same relaxation runs were performed on systems with H_2_O in place of H_2_O_2_.

### 4.4. Umbrella Sampling

Umbrella sampling (US) simulations were performed to quantify the free energy associated with H_2_O_2_ access to the H3C96 and H3C110. Based on the exit pathways observed upon relaxation of the final steered MD conformations, one pathway was simulated for H_2_O_2_ access to H3C96, and two pathways (top and bottom) for access to H3C110. Each pathway was simulated in six independent replicas, starting from conformations observed along the spontaneous exit routes with H_2_O_2_ at ~10 Å from the target cysteine. Because the C110 cavity is symmetric, the bottom route can be entered on either face (through H3D106 or H3′D106); the entry face was assigned for each replica as the D106 copy closest to H_2_O_2_ in the channel region (5–12 Å).

For the H3C96 site, 23 windows were simulated along a collective variable (CV) defined as the distance between the centers of mass of H_2_O_2_ and of the C96 side chain, from 14.5 to 3.5 Å with 0.5 Å steps. As for the H3C110 site, 30 windows were simulated along the distance between the centers of mass of H_2_O_2_ and of both C110 side chains (which face each other at the H3–H3′ interface), so as to describe H_2_O_2_ entering the shared cavity without biasing toward either copy, from 15.0 to 0.5 Å with 0.5 Å steps. Each window was equilibrated for 10 ns, followed by a 50 ns production run, with a force constant of 10 kcal/mol/Å^2^.

The potential of mean force (PMF) of each replica was reconstructed from the production runs with the weighted histogram analysis method (WHAM) [45], using a convergence tolerance of 10^−4^ and 200 bootstrap trials to estimate statistical uncertainties. Each PMF was set to zero on its own bulk plateau (12–15 Å for C96, 11–15 Å for C110). Overlap of the CV distributions of adjacent windows was checked for every replica (Figure S20).

To identify what sets the cost of access, structural descriptors were computed with cpptraj on all production frames and averaged within 0.5 Å bins of the CV. Since the bias acts on the CV only, these conditional averages require no reweighting. Desolvation was monitored as the number of water oxygens within 3.4 Å of any H_2_O_2_ atom. Water access to the target cysteine was quantified, for the frames in which an H_2_O_2_ oxygen is within 3.5 Å of the Sγ atom, there is the presence of at least one water within 3.6 or 5.0 Å of Sγ, and when there is a bridging water, i.e. one whose oxygen lay within 3.6 Å of Sγ and within 3.5 Å of an H_2_O_2_ oxygen. The gate of each route was monitored through the H4F61 χ_1_ dihedral (C96), the H3H113–H3′D123 latch distance on the side of entry (C110, top), and the distance between the H3′I130 Cδ1 atom and the C110 Sγ atom on the entry face (C110, bottom). A gate was considered open when its descriptor left the range sampled with H_2_O_2_ in bulk, i.e. for F61 a χ_1_ outside the *trans* rotamer and for the two distances a value above the 99^th^ percentile of the bulk distribution (3.55 and 5.21 Å, respectively). Pre-organization of the candidate bases was followed as the distance between the carboxylate oxygens of H3E97 (C96) or H3D123 (C110) and the cysteine Hγ atom, a value below 2.5 Å being considered compatible with proton transfer. These descriptors are reported as mean ± standard error over the six replicas.

### 4.5. QM/MM

Given the objective to study the sulfenylation mechanism, which involves bond cleavage and formation, an electronic structure treatment of the reacting species is required. Therefore, we employed the hybrid quantum mechanics/molecular mechanics (QM/MM)[46, 47]molecular dynamics method, treating the reaction active sites quantum mechanically by density functional theory (DFT), while the rest of the biomolecule and solvent are treated with the force field approach. The *ω*B97X [48] functional was used with def2-SVP[49] basis sets for the QM part.

A challenging task is to determine the size of the electronic environment around the central reactive site (the cysteine residue and the H_2_O_2_ molecule). That is, defining the QM region with the appropriate surrounding atoms involved in important charge transfer with the reactive site, within economical computational cost. To mitigate this challenge, we performed the charge shift analysis (CSA) scheme developed by Kulik’s group[50], which is based on the determination of the electronic density redistribution in the absence of the reactive site, thereby, identifying the residues significantly involved in charge transfer with the site. CSA calculations were performed for the systems with targeted sulfenylation at positions C96 and C110. The residues with most charge transfer are given in Table S1. From the results, the C96 protein environment is clearly more hydrophobic than that of C110. With the help of the CSA analysis, consideration of proximity of the residues/atoms to the active site, and some chemical intuition with a balance for simulation cost, the QM region is selected (Figure S21). The link atom approach was applied to treat the QM/MM boundaries, and amino acids sitting at the boundary were “cut” between Cα and Cβ. About 105 to 151 atoms are included in the QM region for the different systems studied.

The QM/MM systems were equilibrated in the NPT ensemble at constant temperature and pressure for 10 ps at the QM/MM level prior to the mechanistic production run. From these dynamics, snapshots of trajectories were selected as starting structures for the mechanistic studies.

### 4.6. QM/MM Umbrella Sampling

Umbrella sampling was performed to explore the reaction mechanism. The hypothesized reaction pathway involves the deprotonation of the cysteine accompanied by the oxidation to sulfenic acid. The deprotonation of the Cys-SH to the more reactive oxidized form (Cys-S^−^) could be either by : i) the H_2_O_2_, route A; ii) a neighbouring glutamate/aspatate, route B; or iii) a neighbouring glutamate/aspatate facilitated by the presence of a water molecule, route C (Figure 3).

In the case of deprotonation by the H_2_O_2_ molecule, the reaction coordinate (RC) was defined by: *RC* = *d*(S_cys_H_cys_) − *d*(H_cys_O_r_), that is, the difference between the Sγ-Hγ distance of the Cys and the Hγ-O_r_ distance (where O_r_ is the reacting oxygen of the peroxide). 11 windows with 0.15 Å step were simulated for H3C96 along the CV from −0.60 to 0.90 Å, while 14 windows were simulated at 0.14 Å step for H3C110 from −0.55 to 1.41 Å. The RC for the deprotonation by glutamate/aspatate on the other hand was defined by: *d*(S_cys_H_cys_) − *d*(H_cys_O_CO−_), where O_CO−_ is the carboxylate reacting oxygen. For the C110, route B was simulated by 13 windows with 0.15 Å step along CV from −0.60 to 1.20 Å, while route C deprotonation was facilitated by the presence of a water molecule in hydrogen-bonding with the Cys-Sγ. The glutamate in the proximity of H3C96 was oriented away from the Cys-Sγ. Therefore, an US was done to reorient it by the rotation about glutamate C_β_-C_γ_ bond as RC in 17 windows at 15° steps from a CV value of −167° to 73°. A further US was conducted for the deprotonation once reoriented, employing the *d*(S_cys_H_cys_) − *d*(H_cys_O_CO−_) CV from 0.08 to 1.33 Å in 11 windows at 0.13 Å step.

The RC for the S-O bond formation was defined by: *d*(O_r_ O_n_) − *d*(O_n_S_cys_), where O_n_ is the non-reacting peroxide oxygen. For the C96, 32 windows were simulated along −3.68 to 2.32 Å CV with 0.25 Å steps. As for C110, 25 and 41 windows were simulated with 0.14 and 0.15 Å steps for routes A and C, respectively, along CV values from −1.32 to 1.20 Å and −3.05 to 2.35 Å.

A 1 ps equilibration followed by 5 ps production run were performed for all windows, with a force constant ranging from 170 to 200 kcal/mol/Å^2^ for the distance CVs and 120 kcal/mol/Å^2^, for the dihedral RCs. Overlap of the CV distributions of adjacent windows were checked for every run (Figure S22). The reconstruction of the PMF was performed by the WHAM method as previously described.

### 4.7. Structural analysis of oxidized nucleosomes

#### DNA opening analysis

To quantify DNA opening along the simulations, dyad-to-end distances were monitored between the center of mass of the DNA backbone atoms of the two central base pairs at the dyad (i.e., residues 73:222 and 74:221) and that of two base pairs at each DNA end (i.e., residues 2:293 and 3:292 at one end, and 145:150 and 146:149 at the other end). Unwrapping was quantified as the percentage of frames in which at least one of the two dyad-to-end distances exceeds 40 Å.

#### DNA flexibility analysis

The flexibility analysis was performed using a PCAbased Python script previously applied to other nucleosomal systems [14, 51, 52]. It uses the inverse distance between each pair of residues as internal coordinates, extracted directly from the MD trajectories. The eigenvectors of the resulting covariance matrix describe the principal modes of motion, and their eigenvalues the amplitude of these motions. The contribution of each residue to these modes then quantifies how it participates in the collective dynamics of the system. Here, the analysis was restricted to the DNA, so as to evaluate the contribution of each nucleotide to DNA flexibility.

#### InterMap interaction analysis

##### Interaction detection

Intermolecular interactions were detected with the InterMap software [21] using its default geometrical definitions. Monitored interaction types included close contacts, van der Waals contacts, hydrogen bonds (donor and acceptor), ionic interactions, cation–*π*, - stacking and hydrophobic contacts. Interactions were analyzed at the residue level: two residues (amino acids, nucleotides or H_2_O_2_) were considered in contact in a given frame if at least one of these interactions was detected between them. Histone residues were numbered according to the histone sequences of PDB 1KX5 [34] in all systems, the two copies of each histone being distinguished by a prime (e.g. H3 and H3′).

##### H_2_O_2_ binding

For the unbiased simulations with free H_2_O_2_, every saved frame was analyzed. An H_2_O_2_ molecule was considered bound to a residue, or to a pocket, when it forms at least one close contact, van der Waals contact or hydrogen bond with that residue (or with any of the pocket residues). Interruptions of not more than 0.1 ns were not counted as unbinding. For each residue and pocket, we report the percentage of simulation time with at least one H_2_O_2_ bound and the longest continuous residence of a single H_2_O_2_ molecule in each replica. The partition of H_2_O_2_ between DNA and histones was obtained by counting, in each frame, the molecules in contact with either partner, and normalized by the number of heavy atoms of each partner. The same definitions were used to identify the residues in contact with H_2_O_2_ along the umbrella sampling pathways.

##### Contact prevalence in the modified nucleosomes

For the simulations of modified and control (unmodified) nucleosomes, one frame every 1 ns was analyzed over the five concatenated replicas (~10,000 frames per system). The prevalence of a residue pair was defined as the percentage of frames in which the two residues are in contact. Only pairs with a prevalence above 30% in at least one system were retained. The effect of each modification was then measured as the change in prevalence (Δ) relative to its own unmodified reference: H3.1 for H3.1soh, and H3.2 for H3.2soh, H3.2sno and H3.2ocs. This analysis was performed separately for the H3–H4, H3–H3′, (H3–H4)_2_–H2A/H2B, (H3–H4)_2_–H2A′/H2B′ and DNA–histone interfaces, restricted to the histone-fold core (H3/H3′ 44–135, H4/H4′ 24–102, H2A/H2A′ 16–119, H2B/H2B′ 34–125). For each interface, the 15 pairs with the largest |Δ| across the four modified systems were selected, together with all contacts involving the modified cysteine. A contact was considered lost when Δ ≤ −25 and gained when Δ ≥ +25.

##### Dimer–tetramer native contacts

Native contacts were defined as the residue pairs with a prevalence of at least 70% in both unmodified H3.1 and H3.2 nucleosomes. For each H2A/H2B dimer, two sets were distinguished: the core contacts with the (H3–H4)_2_ tetramer, formed by the H4–H2B four-helix bundle and the H2A docking domain (residues 80–109), and the contacts of the H2A C-terminal tail (residues 110–118). Tails were excluded from the core set (H3 1–37, H4 1–23, H2A 1–15 and 110–129, H2B 1–30).

#### ComPASS communication pathways analysis

The ComPASS tool was used to identify the communication network of each nucleosome system [23]. Starting from the MD trajectories of a given system, Com-PASS extracts the following properties to build an adjacency matrix: generalized correlations, non-covalent interactions, inter-residue distances and their variance (communication propensity). A graph is then built, where nodes are amino acids and/or nucleotides; edges are defined by applying a distance cutoff to the average minimum distance matrix, and are then weighted using the previously computed adjacency matrix. We then identify the following notions from the graph: *i)* shortest pathways between every pair of residues, *ii)* hotspots (based on centrality measures), *iii)* cliques (fully connected groups of residues), and *iv)* communities (highly interconnected groups of residues that may share a common functional role). Finally, we sort the shortest pathways based on their weight and extract the top 10% for further analysis of communication pathways and allosteric signaling within a given system.

Each modified network was compared with that of its unmodified reference. Path usage was defined, for each edge, as the fraction of top-10% pathways that cross it, and difference maps were built from the change in path usage between the modified and reference systems. For each pair of nucleosome regions, the top pathways connecting them were classified as shared (present in both networks), lost or gained, relative to the percentage of reference pathways kept. The involvement of the H3/H3′ latch was quantified as the percentage of pathways passing within five residues of H3 residues 39–44 in either copy. Finally, to assess the role of the histone tails, the full-length and tailless networks of each system were compared by matching their pathways by endpoint pairs and computing the difference in route length.

#### MDpocket binding pocket analysis

To characterize binding regions close to H3C110 and H3C96, pockets were tracked along the MD ensembles using the MDpocket module of the fpocket software suite with default parameters [53]. Analyses were performed on the control and oxidized systems on concatenated MD replicas, keeping 1 frame out of every 5.

## CRediT authorship contribution statement

**Samuel D. Mador:** Data curation, Formal analysis, Writing - Original draft preparation, Writing - review and editing. **Roy González-Alemán:** Formal analysis, Writing - Original draft preparation, Writing - review and editing. **Yuya Qiu:** Formal analysis, Writing - Original draft preparation, Writing - review and editing. **Francisco Javier Patiño- Lopéz:** Data curation. **Yani Kedjar:** Formal analysis. **Mailys Duch:** Formal analysis. **Yasaman Karami:** Formal analysis, Writing - Original draft preparation, Writing - review and editing. **Emmanuelle Bignon:** Conceptualization, Funding acquisition, Project administration, Supervision, Data curation, Formal analysis, Writing - Original draft preparation, Writing - review and editing.

## Data availability

Data for this article, including MD trajectories of oxidized nucleosomes and input files, are available on Zenodo[54] and on the DynaRepo database [55].

## Acknowledgments

The authors thank Frédéric Beck for his help with data deposition on DynaRepo, and Monha Pepe for fruitful discussions.

## Declaration of generative AI in scientific writing

During the preparation of this work the authors used Claude (Anthropic) in order to improve the language and readability of the manuscript. After using this tool, the authors reviewed and edited the content as needed and take full responsibility for the content of the publication.

