## Supplementary material for "Histone H3 cysteines as redox switches in the nucleosome: molecular modeling of S-sulfenylation and its structural consequences": Supp Fig S1-S21, Supp Table S1

for

---

#### This file includes

- Supplementary Figures S1 to S21
- Supplementary Table S1

---

### Supplementary Figures

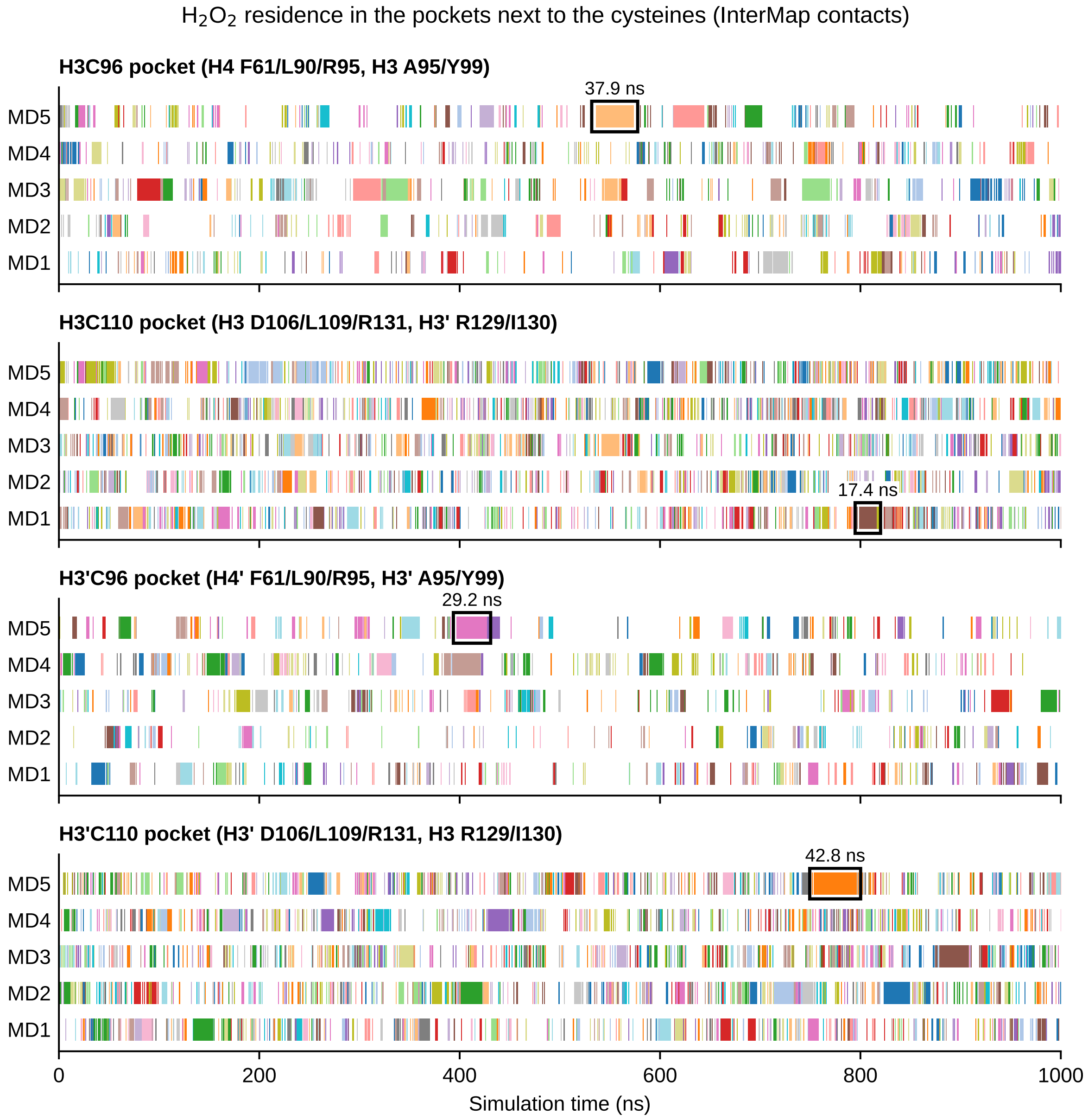


**Figure S1. Time series of $H_2O_2$ residence in the pockets next to C96 and C110 in both H3 copies.** Each bar is one continuous residence event of a single $H_2O_2$ molecule in the pocket, colored by molecule, along the five unbiased MD replicas of the nucleosome core particle. An $H_2O_2$ molecule is considered bound when InterMap detects at least one interaction (close contact, van der Waals contact or hydrogen bond) with a residue of the pocket. Interruptions of up to 0.1 ns are not counted as unbinding. Pockets: H4 F61/L90/R95 and H3 A95/Y99 for C96; H3 D106/L109/R131 and H3′ R129/I130 for C110 (residues swapped between copies for H3′C96 and H3′C110). The longest residence event observed in each pocket is boxed and labeled in black.

‘

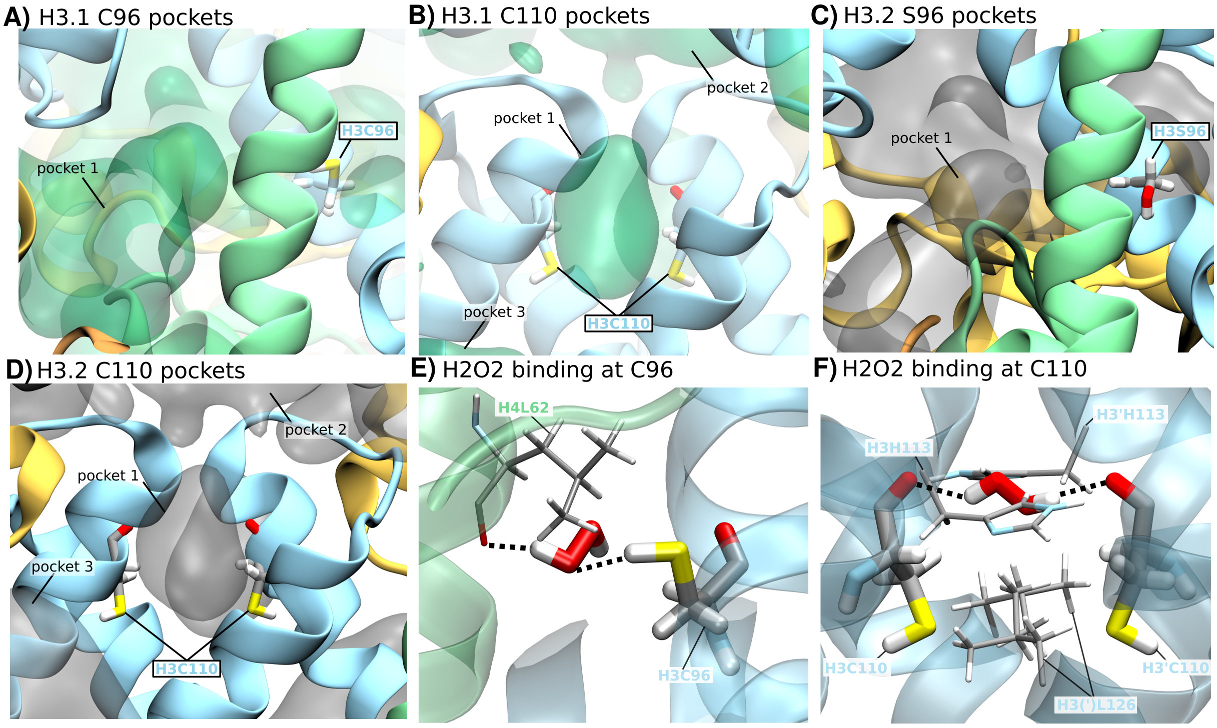


**Figure S2. MDpocket cavity analysis and $H_2O_2$ binding mode at the H3C96 and H3C110 sites.** (A) Pocket identified by MDpocket analysis of MD simulations of the H3.1 nucleosome, in the vicinity of H3C96 (pocket 1). (B) As in (A), for the pockets identified near H3C110/H3′C110 in the H3.1 nucleosome (pockets 1–3). (C) As in (A), for the pocket identified near H3S96 in the unmodified H3.2 control nucleosome (pocket 1), supporting its conservation independently of the C96 mutation. (D) As in (B), for the pockets identified near H3C110/H3′C110 in the H3.2 nucleosome (pockets 1–3). (E) Representative binding mode of $H_2O_2$ within pocket 1 near H3C96, illustrating its elongated conformation simultaneously hydrogen-bonded (dashed lines) to the H4L62 backbone carbonyl and to the H3C96 thiol. (F) As in (E), for $H_2O_2$ bound within pocket 1 near H3C110/H3′C110, hydrogen-bonded to both the H3H113 and H3′H113 side chains.

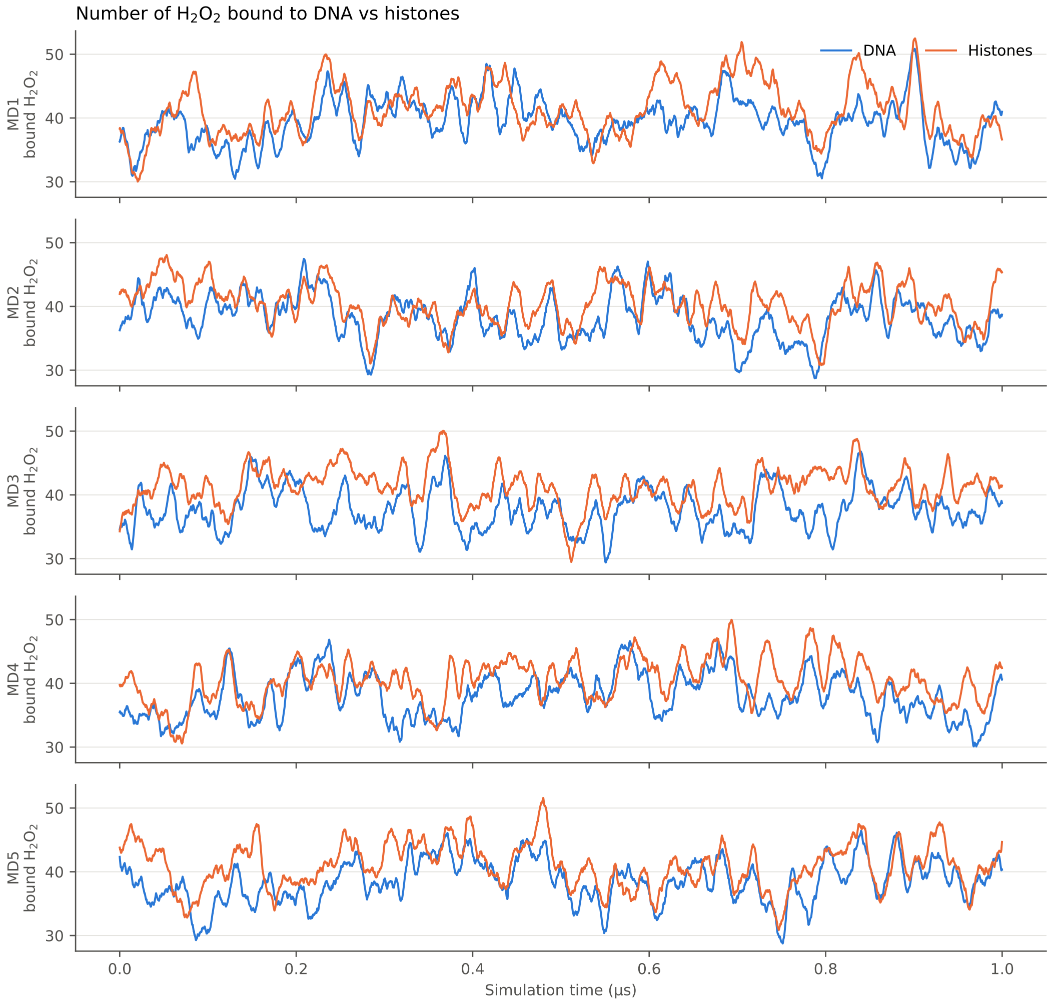


**Figure S3. Number of $H_2O_2$ molecules bound to DNA and histones over time.** Number of $H_2O_2$ molecules bound to DNA (blue) and histones (orange) along each MD replica (10 ns running average). Binding is defined as any InterMap contact (close contact, van der Waals contact or hydrogen bond).

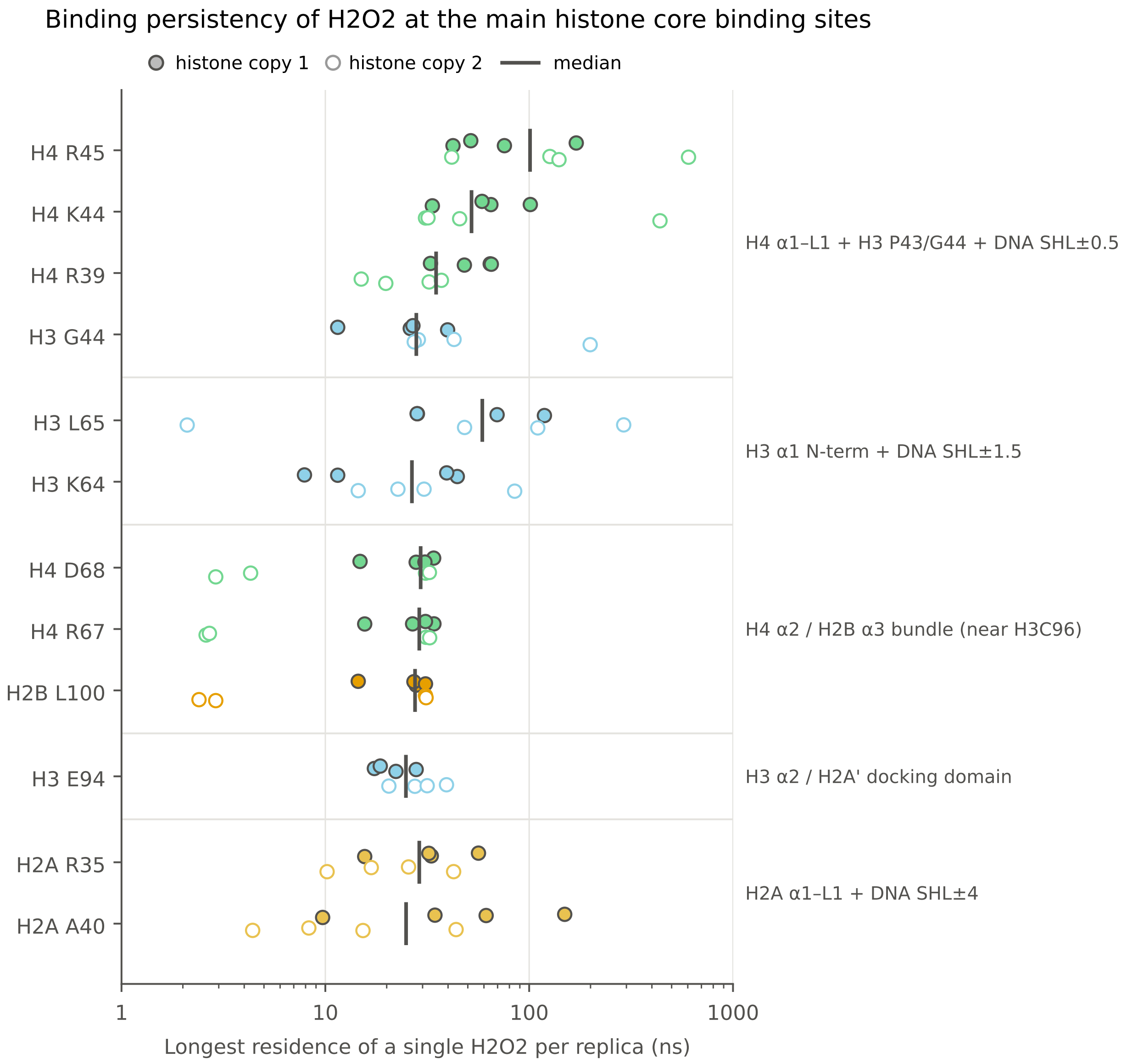


**Figure S4. Binding persistence of $H_2O_2$ at the main histone core binding sites.** Longest residence time of a single $H_2O_2$ molecule per MD replica at each residue, grouped by binding pocket (filled/open circles: first/second histone copy; vertical bars: median). Binding is defined as any InterMap contact (close contact, van der Waals contact or hydrogen bond).

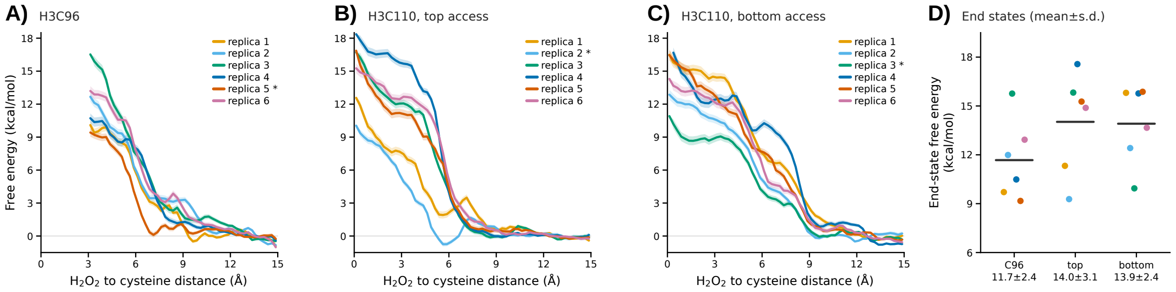


**Figure S5. Umbrella sampling of $H_2O_2$ access - all replicas.** (A–C) Free energy profiles of the six replicas of each route (H3C96; H3C110 top; H3C110 bottom) with WHAM bootstrap error bands. Each profile is zeroed on its own bulk plateau (12–15 Å for C96, 11–15 Å for C110). The lowest-barrier replica, shown in Figure 2–H,I, is marked with an asterisk. (D) End-state free energy of each replica (3–4 Å for C96, 0–1 Å for C110) with the route mean. The replica-to-replica spread (2.4–3.1 kcal/mol) is several times the bootstrap error (0.2–0.5 kcal/mol) and larger than the difference between routes.

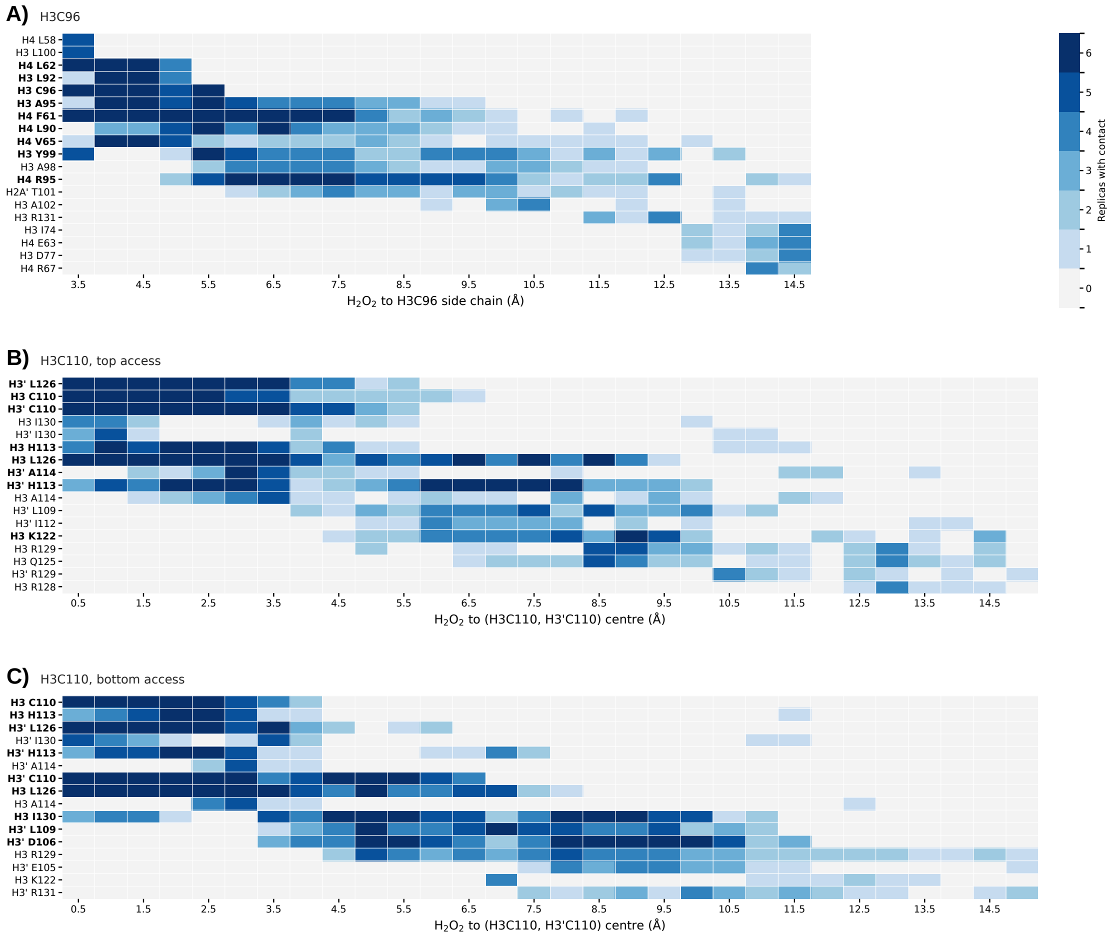


**Figure S6. $H_2O_2$–histone contacts along the umbrella sampling pathways.** Number of replicas (out of six) in which a residue contacts $H_2O_2$ in each umbrella window (considering the 10 last frame of each window), along the pathways to H3C96 (A) and to H3C110 via the top (B) and the bottom (C) of the cavity. Windows run from the bound state (left) to the bulk (right). Only residues contacted in at least four replicas in one window are shown. Bold labels mark residues contacted in all six replicas. For the two symmetric C110 pathways, replicas entering through the other copy of the histones were mirrored onto the majority face.

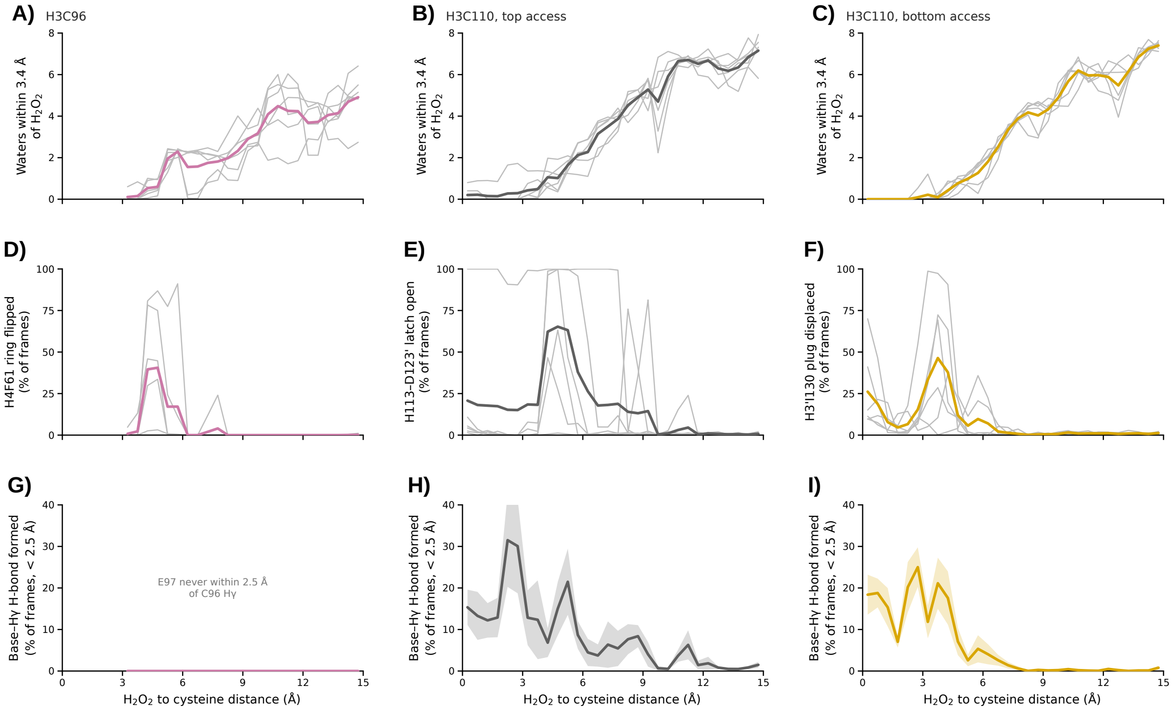


**Figure S7. What the access barrier is made of, route by route.** Columns: H3C96 (A,D,G), H3C110 top (B,E,H) and bottom (C,F,I). Thin grey lines, individual replicas; thick lines, their mean. (A–C) Waters within 3.4 Å of $H_2O_2$. (D–F) Gate open (% of frames): H4F61 $\chi_1$ leaving its bulk rotamer (D), H3H113–H3′D123 latch above the 99th percentile of its bulk distribution (3.55 Å; E), H3′I130 plug above the 99th percentile of its bulk distribution (5.21 Å; F). (G–I) Fraction of frames in which the carboxylate of the candidate base (H3E97 for C96, H3D123 for C110) is within H-bonding distance ($< 2.5$ Å) of the cysteine H$\gamma$. C110 reaches 13–14 % in the bound state on both routes, C96 never.

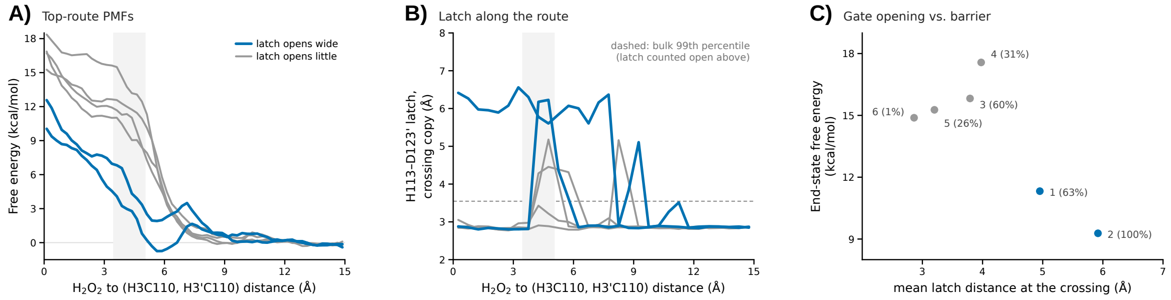


**Figure S8. The top C110 route is bimodal, and the H3H113–H3′D123 latch separates the two modes.** (A) The six top-route profiles, colored by how wide the latch opens while $H_2O_2$ crosses (shaded, 3.5–5 Å). (B) Latch coordinate (the copy $H_2O_2$ passes through) along the reaction coordinate; dashed line, 99th percentile of the bulk distribution, above which the latch is counted as open. (C) End-state free energy against the mean latch distance at the crossing; labels give the replica and the percentage of crossing frames with the latch open. Replicas 1 and 2 (latch opens to 5.4 Å on average) give $10.3 \pm 1.4$ kcal/mol; replicas 3–6 (3.5 Å) give $15.9 \pm 1.2$ kcal/mol.

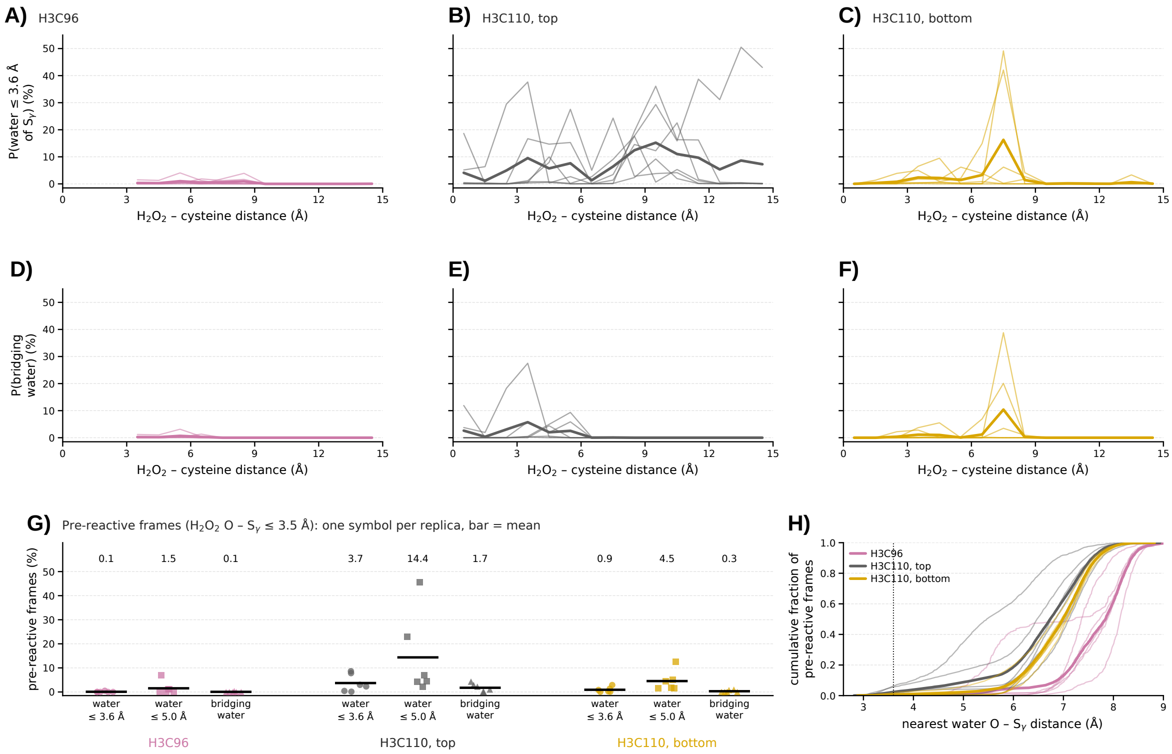


**Figure S9. Water around the target cysteine along the $H_2O_2$ approach.** **(A–C)** Fraction of umbrella-sampling frames in which at least one water lies within 3.6 Å of the Sγ atom of the attacked cysteine (H3C96, A; H3C110 top access, B; H3C110 bottom access, C) as a function of the $H_2O_2$–cysteine distance. For C110, the copy (H3C110 or H3′C110) closest to $H_2O_2$ is analyzed in each frame. **(D–F)** Same for a bridging water, i.e. a water oxygen within 3.6 Å of Sγ and within 3.5 Å of an $H_2O_2$ oxygen in the same frame. **(G)** Pre-reactive frames, defined as $H_2O_2$ O–Sγ $\leq$ 3.5 Å (290–641, 1588–3454 and 3379–4789 frames per replica for C96, C110 top and C110 bottom): fraction with a water within 3.6 Å or 5.0 Å of Sγ and with a bridging water. One symbol per replica; bars and printed values correspond to means. **(H)** Cumulative distribution of the distance between Sγ and the nearest water oxygen in the same pre-reactive frames (thin lines, replicas; thick lines, pooled); the dotted line marks 3.6 Å.

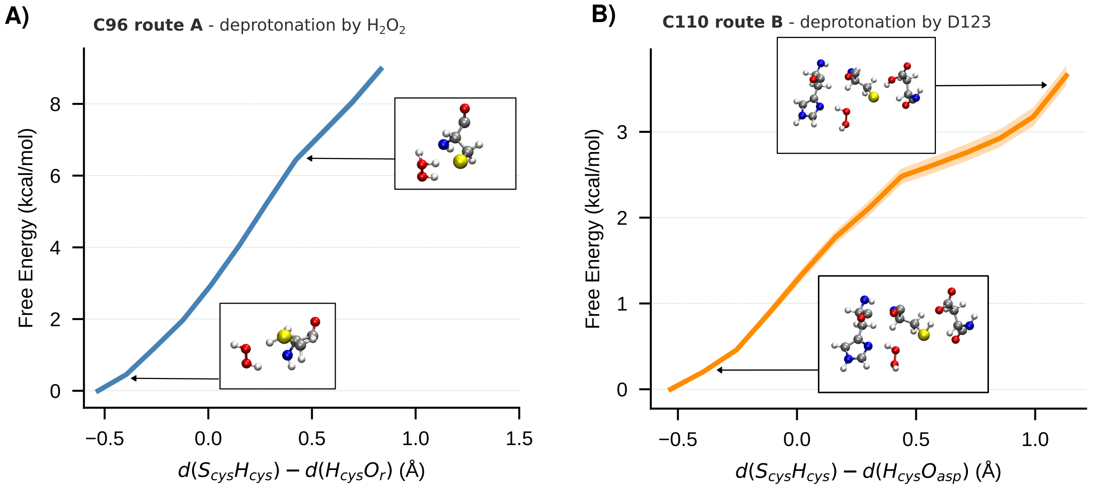


**Figure S10. Free energy profiles for *S*-sulfenylation at C96 and C110.** (A) Free energy profile for the deprotonation of H3C96 by $H_2O_2$ itself (route A). (B) Free energy profile for the deprotonation of H3C110 by H3D123 (route B).

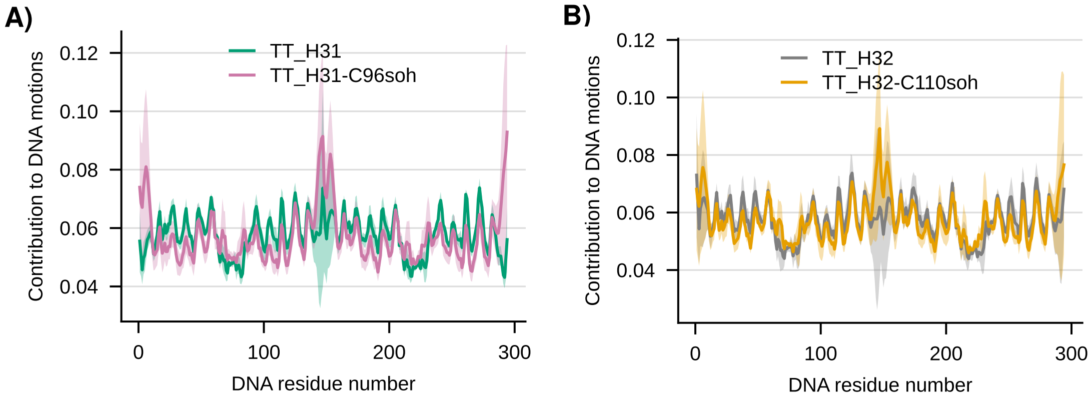


**Figure S11. DNA dynamics of tailless nucleosome core particles bearing cysteine post-translational modifications.** (A) PCA-based per-residue contributions to DNA collective motions during MD simulations of tailless nucleosome core particles containing canonical or $S$-sulfenylated H3.1. (B) As in (A), for canonical or $S$-sulfenylated H3.2.

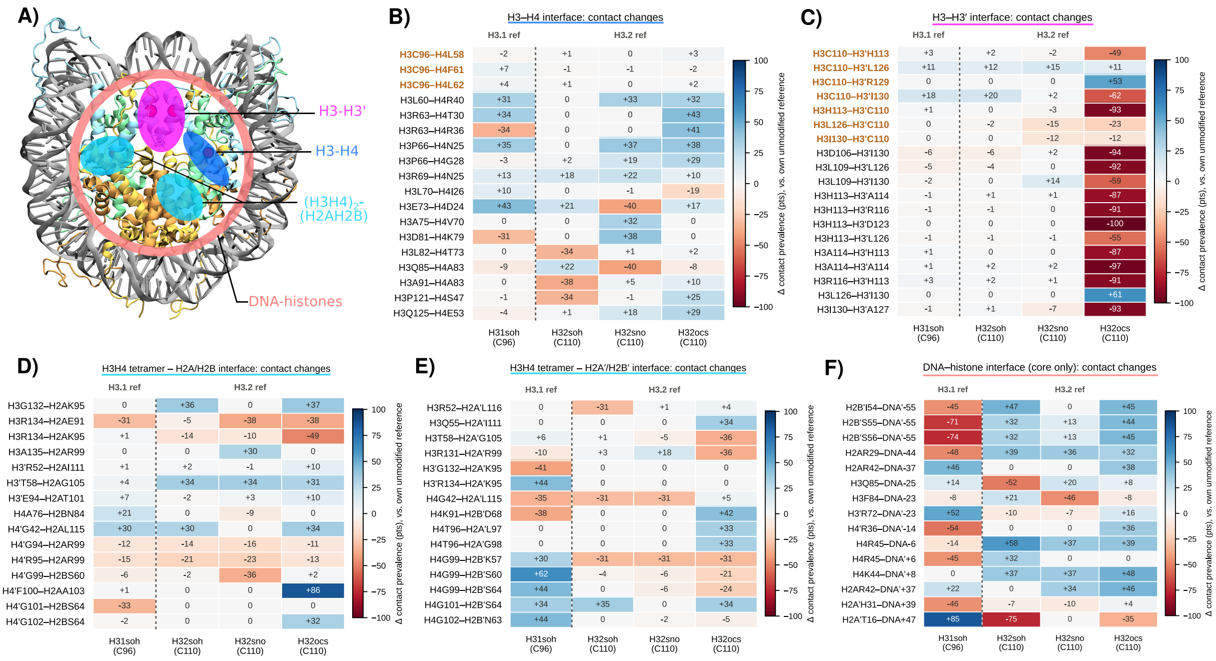


**Figure S12. PTM-induced remodeling of intra-nucleosome contact networks across nucleosome interfaces.** (A) Structural overview of the nucleosome core particle with the interfaces analyzed in (B–F) mapped onto the structure: H3–H3′ (magenta), H3–H4 (dark blue), $(H3H4)_2$–(H2AH2B) tetramer–dimer (cyan), and DNA–histone (salmon). (B–F) Heatmaps of the change in intermolecular contact prevalence (Δ, percentage of MD frames in close contact) for each interface, comparing PTM-bearing systems to their own unmodified reference. Each panel is split into two blocks, separated by a dotted line: left, the H3.1 perturbation upon C96 sulfenylation (H31soh vs. H31); right, the H3.2 perturbation upon C110 $S$-sulfenylation, $S$-nitrosylation, and $S$-sulfonylation (H32soh, H32sno, H32ocs, vs. H32). The top 15 most perturbed interactions are showed for each interface. Red = loss of contact, and blue = gain of contact, with respect to each system's unmodified reference. (B) H3–H4 interface. (C) H3–H3′ interface. (D) $(H3H4)_2$ tetramer–H2A/H2B interface. (E) $(H3H4)_2$ tetramer–H2A′/H2B′ interface. (F) DNA–histone interface. All histone residues in (B–F) are restricted to the histone-fold core (H3/H3′ 44–135, H4/H4′ 24–102, H2A/H2A′ 16–119, H2B/H2B′ 34–125); N-/C-terminal tails are excluded. In (B) and (C), orange bold row labels mark contacts directly involving the modified cysteine (C96 or C110).

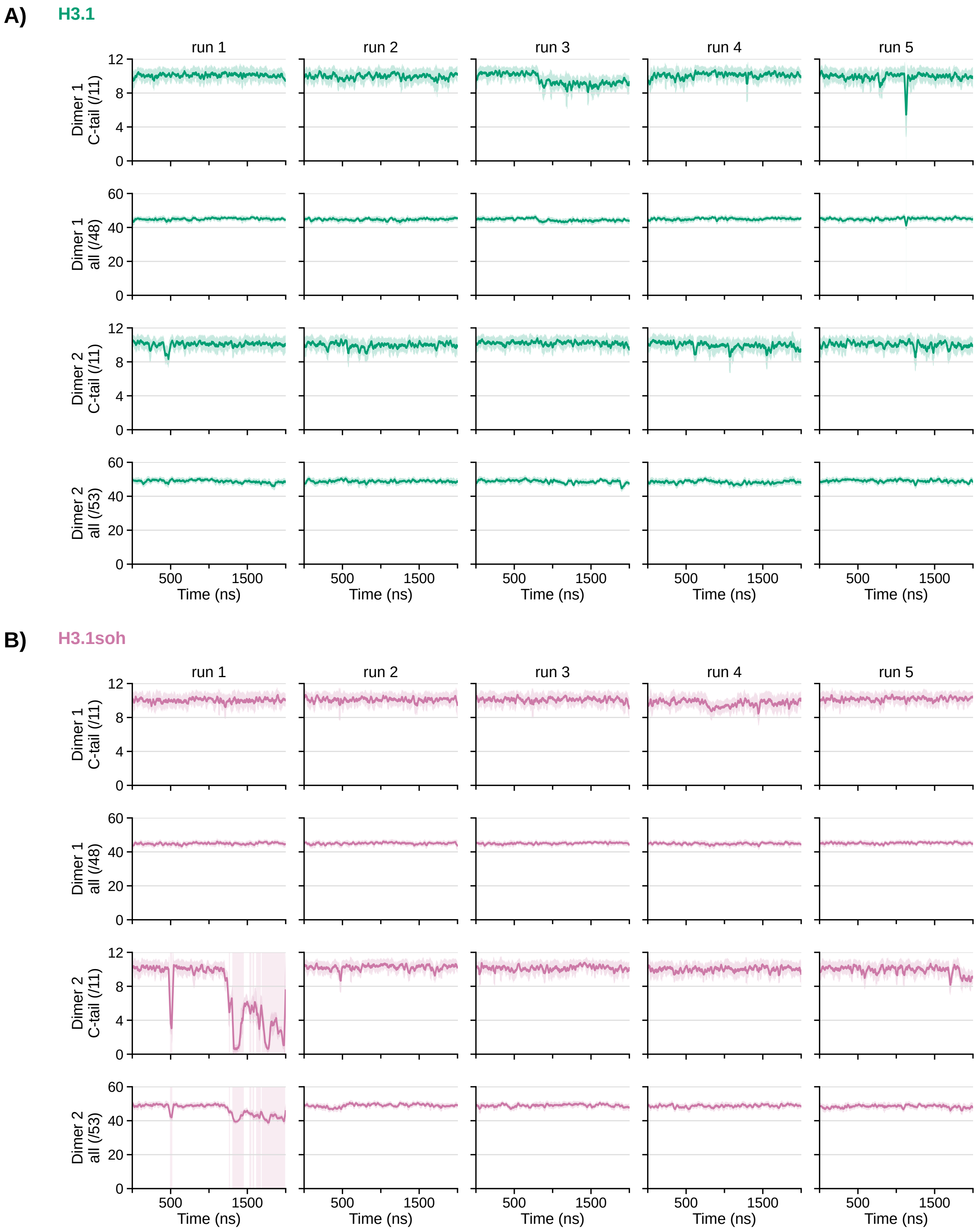


**Figure S13.** Time evolution of the H2A/H2B dimer–$(H3–H4)_2$ tetramer native contacts in the full-length (A) H3.1 and (B) H3.1soh nucleosomes, for each of the five independent 2-$\mu$s replicas (columns). For each H2A/H2B dimer (dimer 1: H2A/H2B; dimer 2: H2A′/H2B′), the first row shows the number of native contacts formed by the H2A C-terminal tail (residues 110–118) with the tetramer, and the second row the total number of dimer–tetramer native contacts. Native contacts are defined as the residue pairs in contact (any InterMap interaction type) for at least 70% of the simulation time in both unmodified full-length references (H3.1 and H3.2); their total number is given in parentheses. Lines show 25-frame running averages and shaded bands the corresponding standard deviations. Shaded backgrounds highlight the frames in which the H2A C-terminal tail retains fewer than half of its native contacts (tail release).

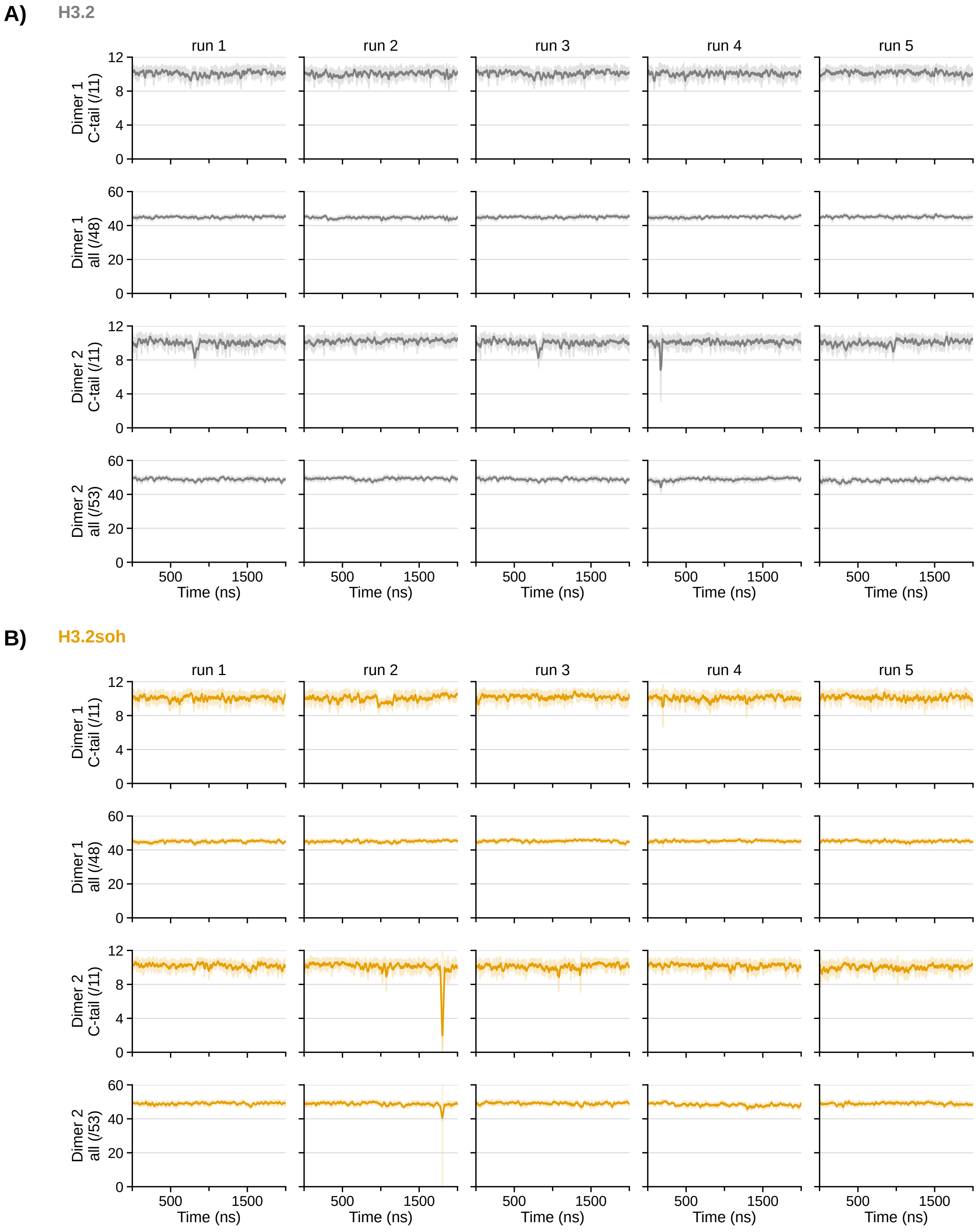


**Figure S14.** Time evolution of the H2A/H2B dimer–$(H3–H4)_2$ tetramer native contacts in the full-length (A) H3.2 and (B) H3.2soh nucleosomes, for each of the five independent 2-$\mu$s replicas. Rows, contact definitions and shading as in Figure S13.

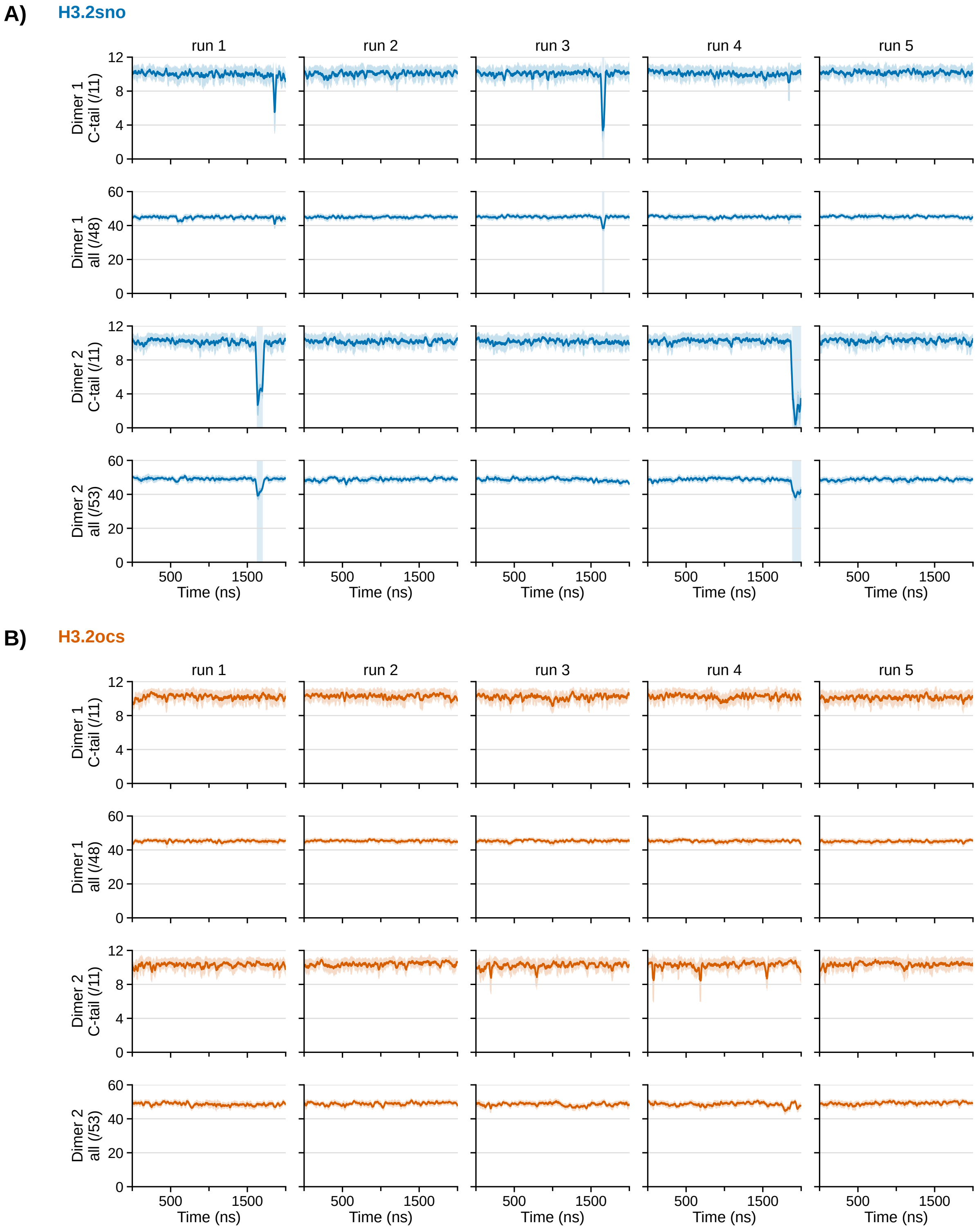


**Figure S15.** Time evolution of the H2A/H2B dimer–(H3–H4)$_2$ tetramer native contacts in the full-length (A) H3.2sno and (B) H3.2ocs nucleosomes, for each of the five independent 2-$\mu$s replicas. Rows, contact definitions and shading as in Figure S13.

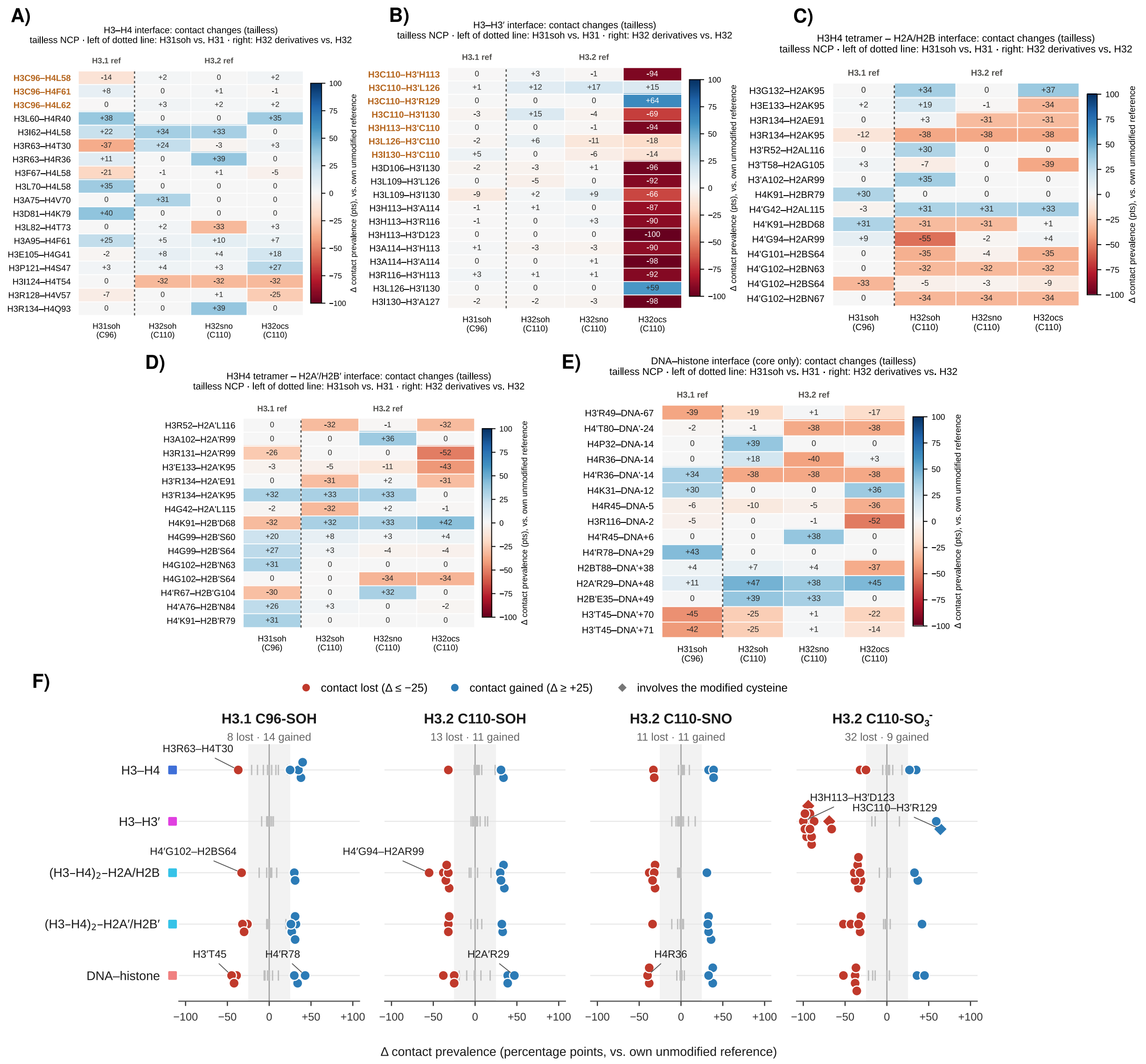


**Figure S16. Redox modification-induced changes in histone–histone and histone–DNA contacts in the tailless nucleosome core particle. (A–E)** Heatmaps of the change in contact prevalence (Δ contact prevalence, percentage points) for the residue pairs most affected by the modifications at each interface, restricted to the histone-fold core. Each modified system is compared to its own unmodified reference: H3.1soh to H3.1 (left of the dotted line), and the H3.2 derivatives (H3.2soh, H3.2sno, H3.2ocs) to H3.2 (right of the dotted line). Blue and red shading indicate a gain or a loss of contact prevalence upon modification, respectively. Residue pairs directly involving the modified cysteine (C96 or C110) are highlighted in orange. **(A)** H3–H4 interface. **(B)** H3–H3′ interface. **(C)** $(H3–H4)_2$ tetramer–H2A/H2B interface. **(D)** $(H3–H4)_2$ tetramer–H2A′/H2B′ interface. **(E)** Histone core–DNA interface. In all panels, columns correspond to H3.1soh (*S*-sulfenylation at C96), H3.2soh (*S*-sulfenylation at C110), H3.2sno (*S*-nitrosylation at C110) and H3.2ocs (over-oxidation of C110 to the cysteine sulfonic acid). **(F)** Summary of panels A–E for each modified system. Each dot is a residue pair whose contact prevalence is lost ($\Delta \leq$ -25, red) or gained ($\Delta \geq +25$, blue) upon modification, one row per interface (colored squares, matching the interfaces of the structure in Figure 6A). Grey ticks indicate pairs with $|\Delta| < 25$, diamonds indicate pairs involving the modified cysteine, and the numbers of contacts lost and gained are given under each system name. A few representative contacts are labeled.

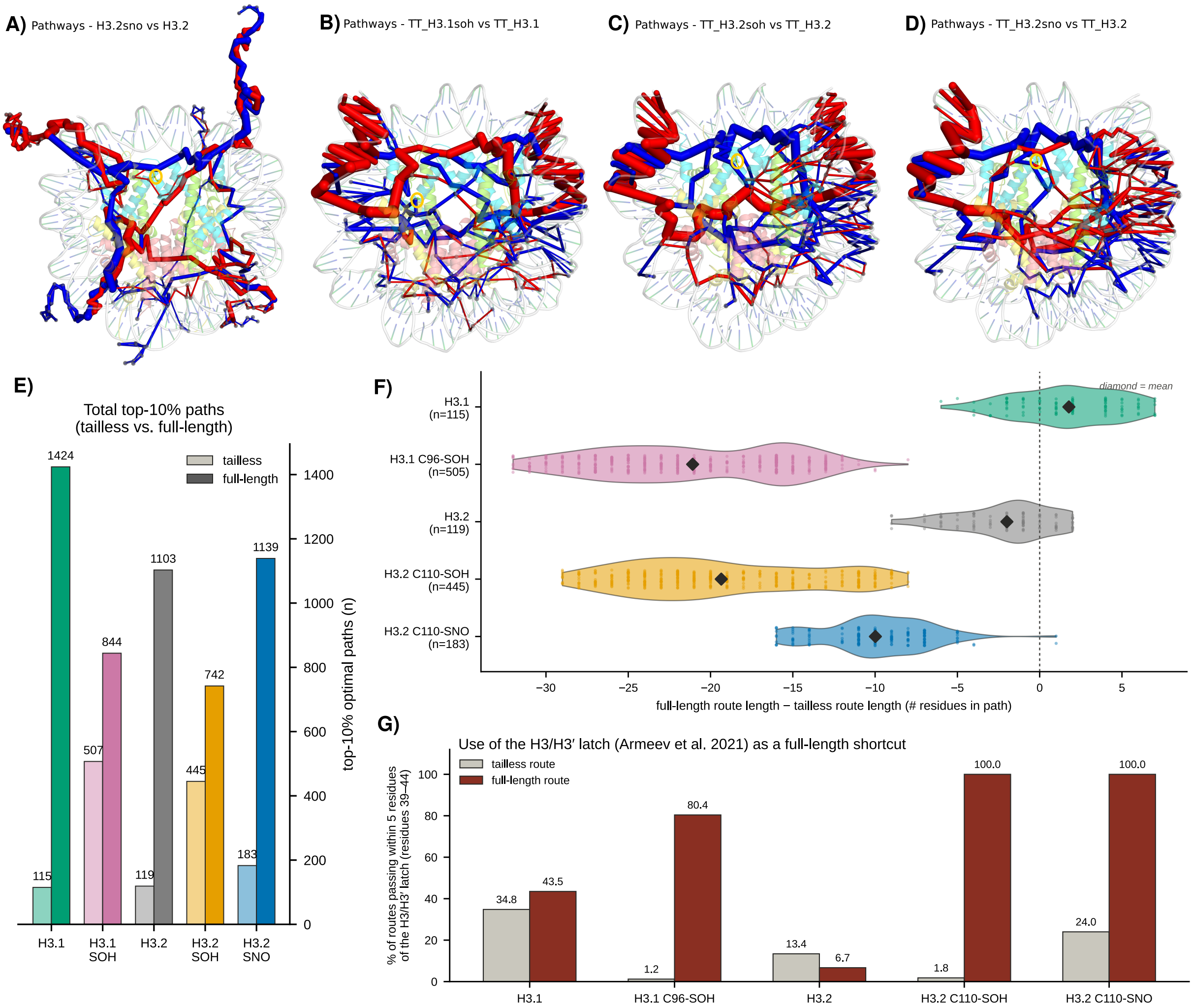


**Figure S17. Cysteine oxidation reroutes the nucleosome communication network, and the histone tails supply the shortcut. (A–D)** Difference maps of the ComPASS optimal-communication network. Each stick is an edge of the top-10% shortest-path network, drawn between consecutive residues along a path (C$\alpha$ for protein, C5′ for DNA; nodes shown as grey spheres). Red marks edges carried by a larger fraction of the top-10% paths in the modified system than in its own unmodified reference, blue a smaller fraction, and stick width scales with the size of that change. Histones are shown as transparent cartoons coloured by type and DNA in pale grey; the modified cysteine is yellow. **(A)** full-length H3.2 C110-SNO versus H3.2. **(B)** tailless H3.1 C96-SOH versus tailless H3.1. **(C)** tailless H3.2 C110-SOH versus tailless H3.2. **(D)** tailless H3.2 C110-SNO versus tailless H3.2. **(E)** Number of optimal paths falling in the top 10% of the shortest-path distribution, for each system in the tailless (light) and full-length (solid) construct. **(F)** For every endpoint pair found in a tailless top-10% path, the length of the corresponding full-length route minus that of the tailless route, in residues; $n$ is the number of endpoint pairs compared. Dots are individual pairs, diamonds the means, and the dashed line marks no change. **(G)** Percentage of those routes passing within five residues of the H3/H3′ latch (residues 39–44), for the tailless route (grey) and the full-length route (dark red) of the same endpoint pairs.

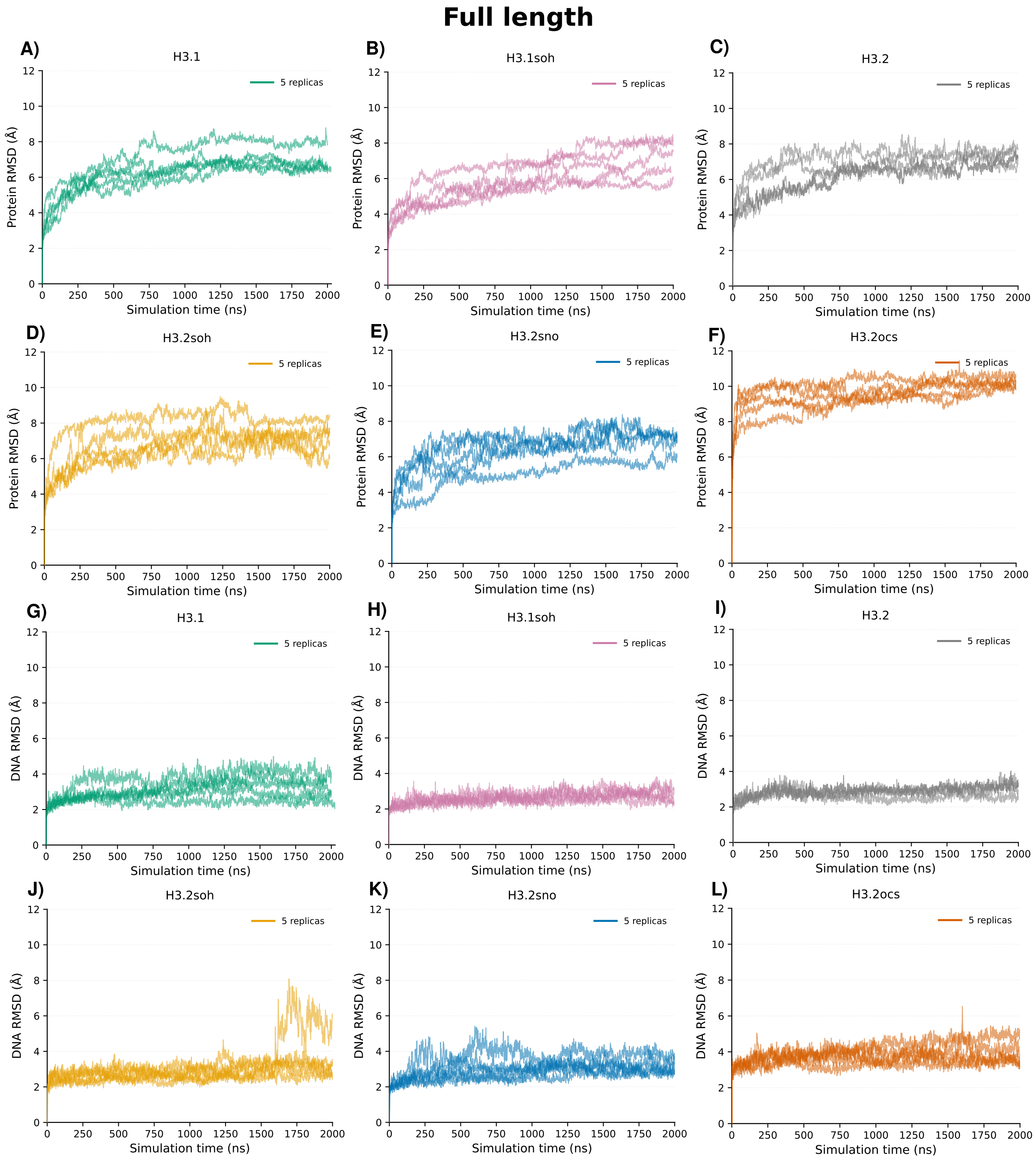


**Figure S18. RMSD monitoring of the full-length nucleosome core particle systems.** Time evolution of the protein backbone RMSD (A–E) and DNA backbone RMSD (F–J) along the five replicas (2 $\mu$s each) of each full-length nucleosome system: H3.1 (A, G), H3.1soh (B, H), H3.2 (C, I), H3.2soh (D, J), H3.2sno (E, K), and H3.2ocs (F, L). RMSD values were computed relative to the starting structure of each replica, following superposition on the corresponding (protein or DNA) atoms.

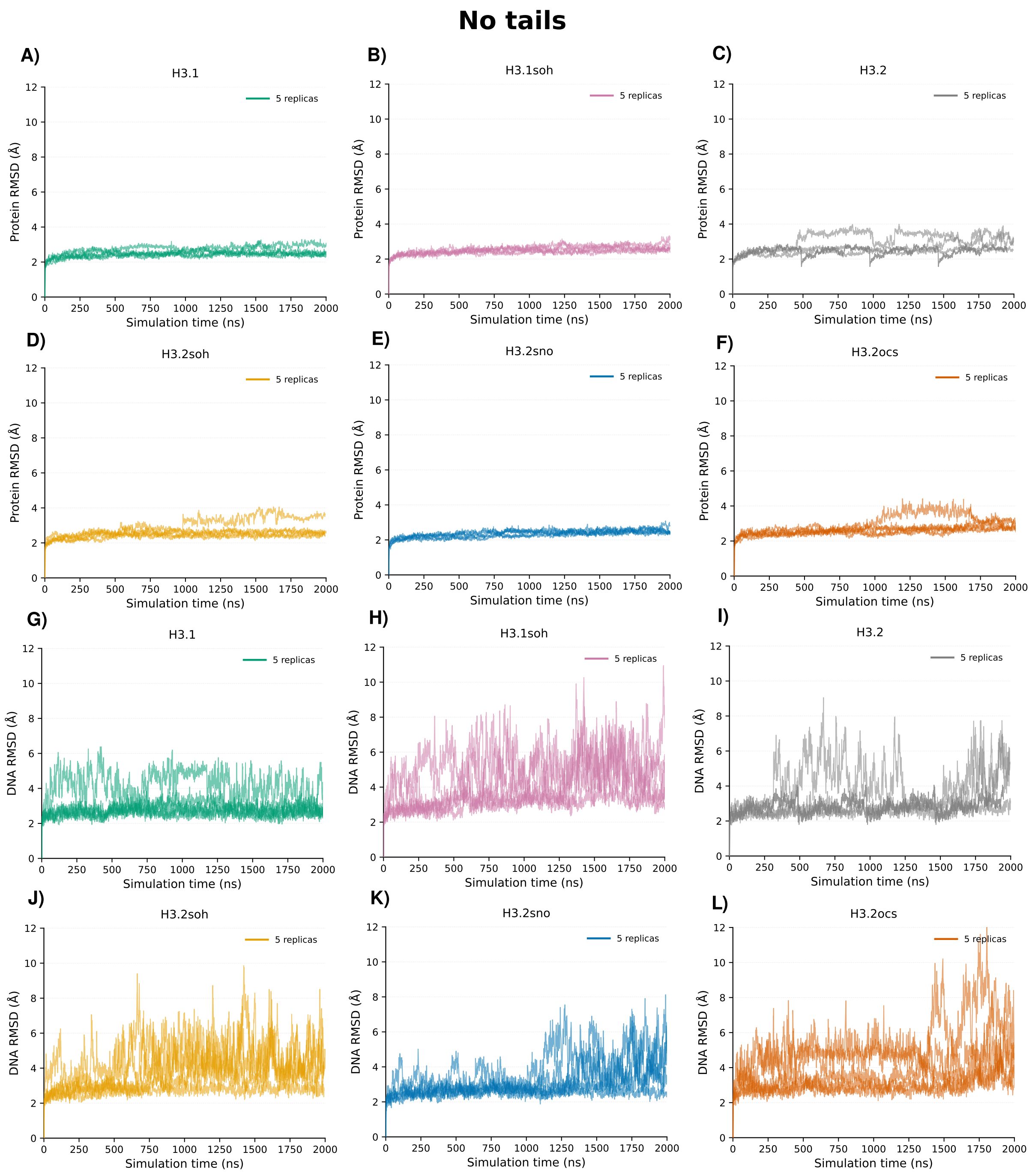


**Figure S19. RMSD monitoring of the tailless nucleosome core particle systems.** Time evolution of the protein backbone RMSD (A–E) and DNA backbone RMSD (F–J) along the five replicas (2 $\mu$s each) of each tailless nucleosome system: H3.1 (A, G), H3.1soh (B, H), H3.2 (C, I), H3.2soh (D, J), H3.2sno (E, K), and H3.2ocs (F, L). RMSD values were computed relative to the starting structure of each replica, following superposition on the corresponding (protein or DNA) atoms.

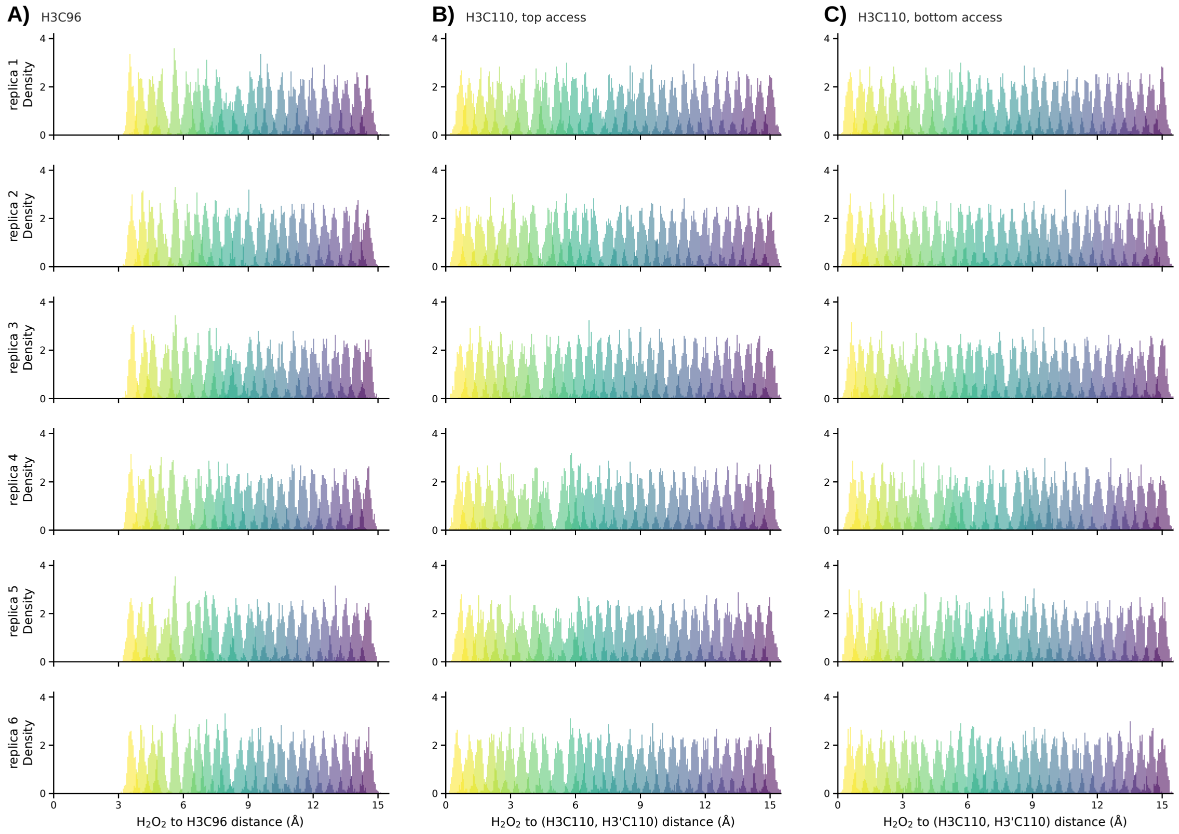


**Figure S20. Overlap of the umbrella sampling windows for all replicas.** Distribution of the $H_2O_2$–cysteine distance sampled in each umbrella window (colored from bulk, purple, to the bound state, yellow) for the six independent replicas (rows) of the access pathway to H3C96 (A; 23 windows per replica), and to H3C110 via the top (B) and the bottom (C) of the cavity (30 windows per replica). Windows are spaced by 0.5 Å with a force constant of 20 kcal/mol/Å$^2$ in the WHAM analysis. Neighboring windows overlap in all replicas (median overlap coefficient 0.14–0.15, minimum 0.01–0.02), the weakest links lying between 4 and 6 Å, where the profile is steepest.

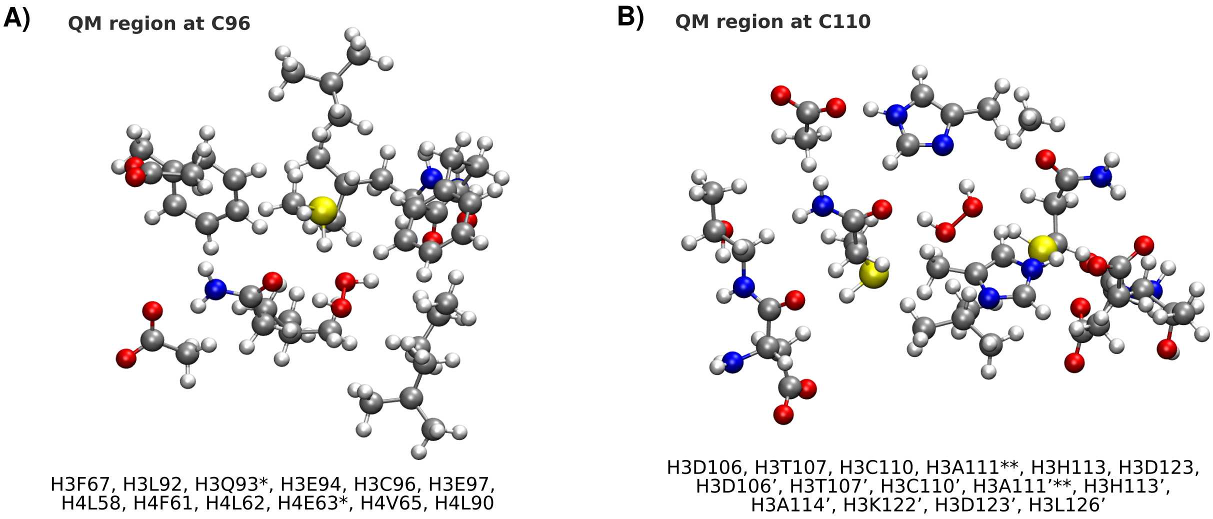


**Figure S21. Composition of the QM region for *S*-sulfenylation at C96 and C110.** Residues included in the QM region for the QM/MM calculations at (A) C96 and (B) C110.

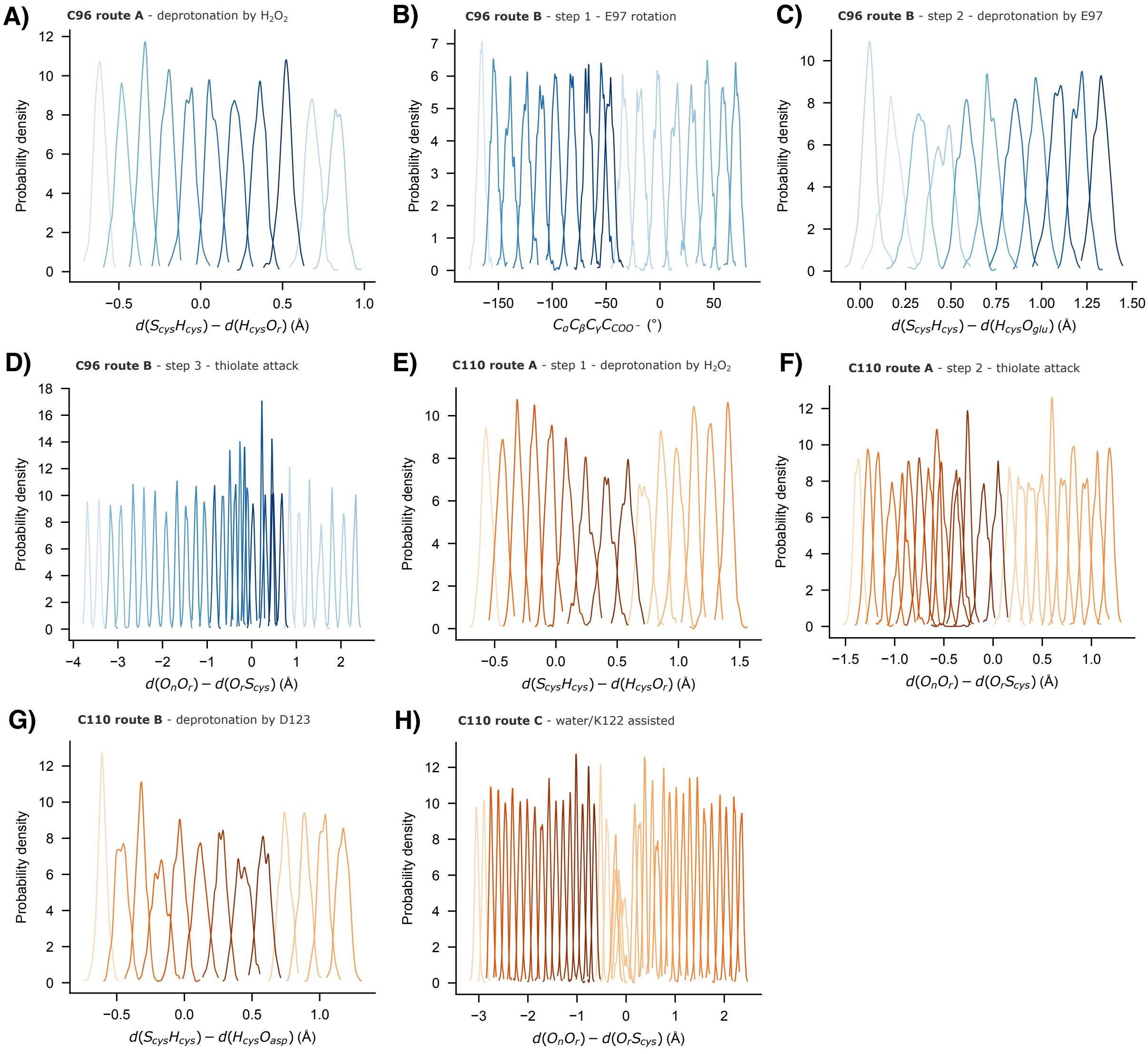


**Figure S22. Overlap of the umbrella sampling windows for all reactivity routes.** Distribution of the CVs sampled in each umbrella window for reactivity at C96 following route A (A), route B (A-D), and for reaction at C110 following route A (E), route B (F-G) and route C (H).

### Supplementary Table

**Table S1. Charge-shift analysis (CSA) of residues surrounding the reactive H3C96 and H3C110 sites.** Change in partial atomic charge ($\Delta q$, $e$) of the reactive site (cysteine + $H_2O_2$) upon computational alanine mutation of each surrounding residue, identified with the CSA scheme of Kulik and coworkers and used together with residue/atom proximity to guide selection of the QM region (Figure S21).

| C96 | | C110 | |
|---|---|---|---|
| Residue | $\Delta q$ | Residue | $\Delta q$ |
| H3F76 | +0.00 | H3D106 | −0.18 |
| H3L92 | +0.01 | H3T107 | +0.27 |
| H3Q93 | −0.02 | H3L109 | +0.08 |
| H3E94 | −0.01 | H3H113 | −0.15 |
| H3A95 | +0.00 | H3A114 | −0.37 |
| H3E97 | +0.02 | H3L126 | −0.44 |
| H3A98 | +0.02 | H3I130 | −0.36 |
| H3Y99 | +0.02 | H3'D106 | −0.40 |
| H3L100 | −0.04 | H3'T107 | +0.22 |
| H4V57 | −0.00 | H3'N108 | +0.15 |
| H4L58 | −0.04 | H3'L109 | −0.15 |
| H4F61 | −0.02 | H3'A111 | −0.06 |
| H4L62 | −0.02 | H3'I112 | −0.18 |
| | | H3'H113 | +0.39 |
| | | H3'D123 | +0.43 |
